# Ultraviolet light-induced collagen degradation inhibits melanoma invasion

**DOI:** 10.1101/2021.01.27.428482

**Authors:** Timothy Budden, Caroline Gaudy, Andrew Porter, Emily Kay, Shilpa Gurung, Charles Earnshaw, Katharina Roeck, Sarah Craig, Víctor Traves, Jean Krutmann, Patricia Muller, Luisa Motta, Sara Zanivan, Angeliki Malliri, Simon J Furney, Eduardo Nagore, Amaya Virós

## Abstract

Ultraviolet radiation (UVR) increases the incidence of cutaneous melanoma^1–4^. The ageing, sun-exposed dermis accumulates UVR damage^5^, and older patients develop more melanomas at UVR-exposed sites^4,6,7^. As fibroblasts are functionally heterogeneous and play key roles in the stromal contribution to cancer^8,9^, we asked whether UVR modifies dermal fibroblast function. Here we confirmed the expression of collagen-cleaving matrix metalloprotein-1 (*MMP1*) by UVR-damaged fibroblasts was persistently upregulated to reduce local levels of collagen 1 (*COL1A1*), and found dermal COL1A1 degradation by MMP1 decreased melanoma invasion. Conversely, we show inhibiting extracellular matrix degradation and MMP1 expression restored melanoma invasion to UVR damaged dermis. We confirmed *in vitro* findings in a cohort of primary cutaneous melanomas of aged humans, showing more cancer cells invade as single cells at the invasive front of melanomas expressing and depositing more collagen. We found collagen and single melanoma cell invasion are robust predictors of poor melanoma-specific survival. These data indicate melanomas arising over UVR-damaged, collagen-poor skin of the elderly are less invasive, and this reduced invasion improves survival. Consequently, although UVR increases tumour incidence, it delays primary melanoma invasion by degrading collagen. However, we show melanoma-associated fibroblasts can restore invasion in low-collagen primary tumours by increasing collagen synthesis. Finally, we demonstrate high *COL1A1* gene expression is a biomarker of poor outcome across a broad range of primary cancers.

## Main

UVR is the major environmental risk factor for the development of melanoma^1,2,10^ and sun exposure is the main cause of rising disease incidence^3^. Whilst the association between UVR and melanoma incidence is well established^10^, there are controversial clinical studies associating sun exposure, or sun damage to the dermis, with improved melanoma survival^11–13^. However, other studies have found no association between sun damage and outcome, and clinical studies show melanomas arising on the scalp and neck, areas likely chronically sun damaged, are linked to poor outcome^14–16^.

UVR damage accumulates with increasing decades of life, and aged patients have worse melanoma survival^17,18^. Therefore, it is possible that chronic UVR damage may lead to shorter melanoma specific survival. However, in common with some non-hormonal cancers, the incidence and mortality of melanoma sharply rise after age 60, and then significantly decrease after age 85^19,20^. suggesting the relationship between cumulative UVR exposure, cutaneous damage, age and melanoma death is not linear.

Previous studies have shown collagen quantity in the extracellular matrix (ECM) modifies melanoma cell behaviour^21^. Surprisingly, both increased^22^ and decreased^23^ deposition of collagen have been linked to malignant behaviour, suggesting the effect of collagen on cancer behaviour extends beyond protein level and scaffold function. In this study we explore how collagen levels in the dermis, which vary according to sun damage and age, affect melanoma survival.

### Somatic mutation burden in dermal fibroblasts correlates with extracellular matrix degradation and collagenase expression

The pivotal task of dermal fibroblasts is to regulate ECM remodelling, including the turnover of collagen^24^. In chronically UVR-damaged skin there is an increase of degraded collagen that is not compensated by new collagen synthesis, contributing to overall ECM degradation. We analysed gene expression in human adult fibroblasts to compare matched UVR-damaged and UVR-protected dermis from healthy donors (median age 42, range 19-66^25^). We used COSMIC total Signature 7 mutations^26^, which indicate UVR-induced damage, as a surrogate marker of accumulated UVR exposure. Strikingly, the most significantly differentially expressed pathway was the ECM pathway (Fig. 1a, b; Extended Data Table 1, 2) demonstrating progressive down-regulation of collagen genes (Extended Data Table 3) and upregulation of matrix metalloproteinases (MMP), including matrix metalloproteinase-1 (*MMP1*), in UVR-damaged adult donor fibroblasts, with increasing signature 7 mutations. Furthermore, *MMP1* was highly expressed in donor fibroblasts from UVR-exposed calves of adults in the Genotype-Tissue Expression cohort^27^ (median TPM =51.85, n =504).

**Figure 1.**
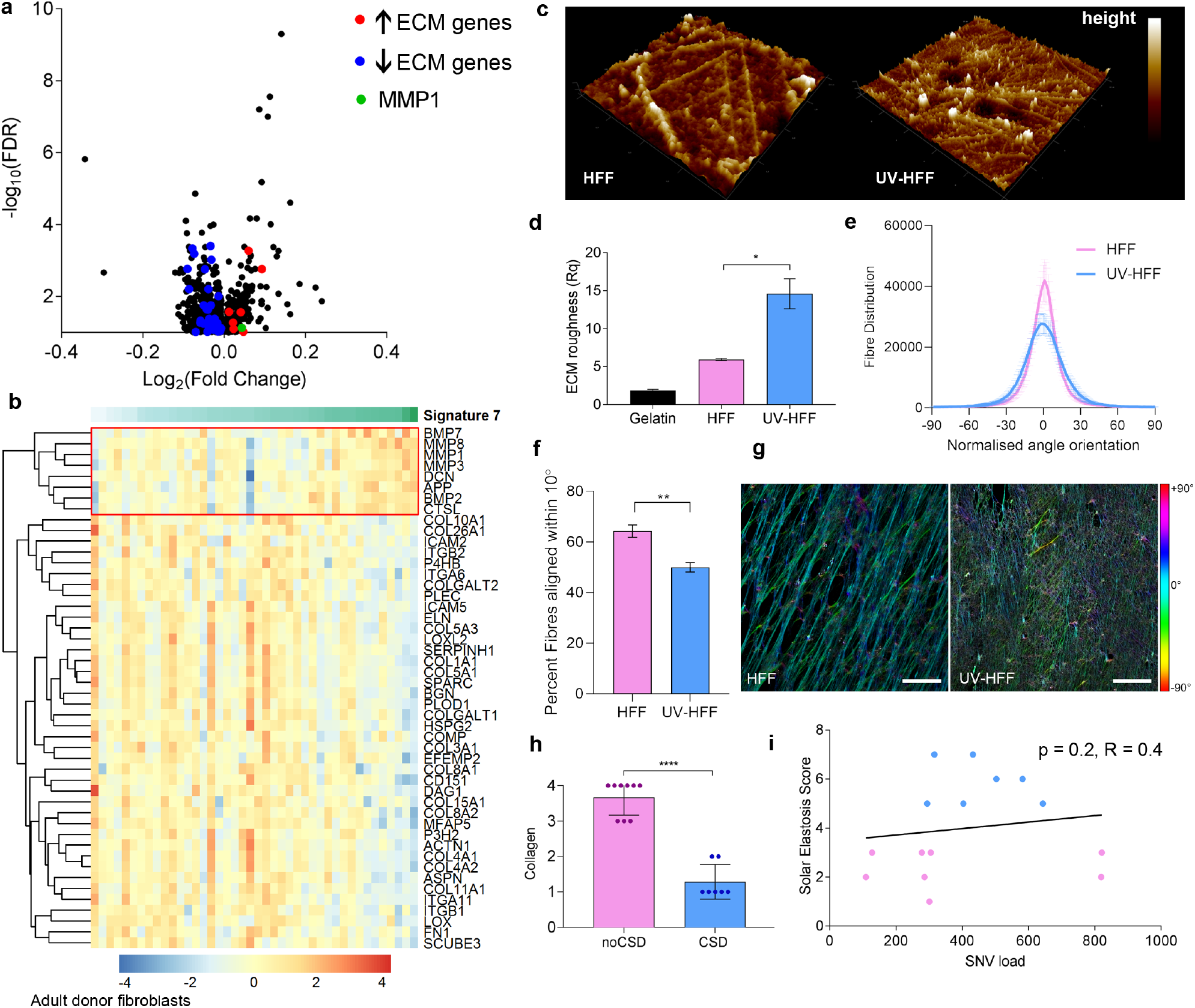
UVR-driven mutations in adult donor dermal fibroblasts correlate with ECM degradation and collagenase expression. **(a)** Volcano plot gene expression and **(b)** extracellular matrix (ECM) heatmap by COSMIC Signature 7 mutations in skin fibroblasts^25^ red: upregulated, blue downregulated genes. **(c)** Atomic force microscopy and **(d)** matrix roughness (Rq) of *in vitro* ECM of HFF and UV-HFF, Mann-Whitney U *p<0.05. **(e)** Quantification of fibre alignment distribution in human foreskin fibroblasts HFF and UV-HFF derived ECM by fibronectin immunofluorescence **(f)** Fraction of fibres within 10° of mode orientation, Mann-Whitney U **p<0.01 **(g)** Immunofluorescence of fibronectin fibres in decellularised HFF and UV-HFF derived ECM, colour coded for orientation of fibre, cyan represents mode, scale bar: 25 μm. **(h)** Masson’s trichrome collagen stain of sun protected (n=9, noCSD) and sun damaged (n=7, CSD) dermis, Mann-Whitney U ****p<0.0001. **(i)** Correlation between SNV load and solar elastosis in fibroblasts, blue: CSD, pink: noCSD, Spearman correlation R =0.40, p =0.2. Error bars: standard error of the mean (bar).

MMP1 cleaves collagen 1 (COL1A1) after acute UVR exposure^28,29^, so we compared the secretion of MMP1, *COL1A1* expression, and ECM collagen deposition of isogenic UVR-naïve fibroblasts from a human fibroblast foreskin cell line (HFF) and UVR-damaged fibroblasts (UV-HFF) 2 weeks after UVR exposure (Extended Data Fig. 1a). We found UV-HFF fibroblasts increased the secretion of MMP1 (Extended Data Fig. 1b) with no compensatory increase in *COL1A1* transcription (Extended Data Fig. 1c) or deposition in the ECM of UV-HFF compared to UVR-naïve fibroblasts (COL1A1 Mean LFQ intensity HFF =33.20, UV-HFF =33.47, q-value =0.16, COL1A2 HFF =32.57, UV-HFF =32.87, q-value =0.08, Extended Data Fig. 1d). Additionally, atomic force microscopy (AFM) topographic imaging suggested UV-HFF fibroblast-generated ECM presented more fragmented, sparser and disorganised matrix fibrils than UVR-naïve, HFF fibroblasts. The higher roughness (Rq) value indicates less symmetry across the ECM surface plane, in keeping with degradation of UV-HFF fibroblast-generated ECM^30–32^ (Fig. 1c, d). Furthermore, immunofluorescent staining of fibronectin fibres in HFF and UV-HFF derived ECM, confirmed that UV-HFF matrices were significantly more disorganised with fewer aligned fibres than matrices generated by HFF fibroblasts (Fig. 1e-g).

Since UVR damage alters fibroblast function, compromising ECM renewal, we compared the density of collagen fibres in chronically sun damaged and sun-protected healthy skin of aged patients (n=16, age >59); confirming reduction of collagen in UVR-damaged dermis^33^ (Fig. 1h). Additionally, we confirmed fibroblasts from tumour-adjacent sun-damaged patient dermis (solar elastosis^33,34^) have higher total somatic mutation burden^5^ (n=13, Fig. 1i), indicating that cumulative UVR leads to dermal ECM degradation and decreased collagen.

### Low collagen concentration and reduced collagen integrity decrease melanoma invasion

To study if UVR damage to fibroblasts driving collagen degradation affects melanoma progression, we compared melanoma invasion in spheroids embedded in matrices of increasing collagen concentrations. Melanoma cell lines present varying degrees of invasion, so we used three cell lines established from tumours bearing different UVR mutation signatures, reflecting UVR and non-UVR tumour origins^35^ (Extended Data Fig. 2a). We found that regardless of the UVR history of the melanoma cell line, the invasion of the three melanoma lines was optimal in 1.5mg/ml collagen, and higher (2.5mg/ml, p <0.0001) and lower collagen concentrations significantly reduced invasion into the ECM (0.25mg/ml, p<0.0001, 0.5mg/ml, p=0.03; Fig. 2a, b, Extended Data Fig. 2b). Additionally, we quantified the number of melanoma cells detaching from the spheroid and invading as single cells, and found single cell invasion optimal within a range of collagen concentrations, decreasing with higher and lower collagen densities (Fig. 2c, d). We generated organotypic dermal constructs with HFF or UV-HFF foreskin human fibroblasts, seeded with melanoma cell lines (Fig. 2e, Extended Data Fig. 2c), and confirmed UV-HFF constructs presented fewer melanoma cells detaching from the tumour edge, singly advancing in the dermis (Sk-mel-28 p=0.04; Fig. 2f). UV-HFF constructs replicated the cardinal features of UVR damage^36–38^, with significantly reduced collagen levels compared to HFF constructs (p<0.0001, Extended Data Fig. 2d, e). Additionally, UV-HFF constructs presented reduced fibronectin (Extended Data Fig. 2f), and no difference in elastin expression compared to HFF constructs (Extended Data Fig. 2g). These data indicate that melanoma invasion is optimal within a range of collagen concentrations. Critically, lower collagen concentrations limit melanoma invasion.

**Figure 2.**
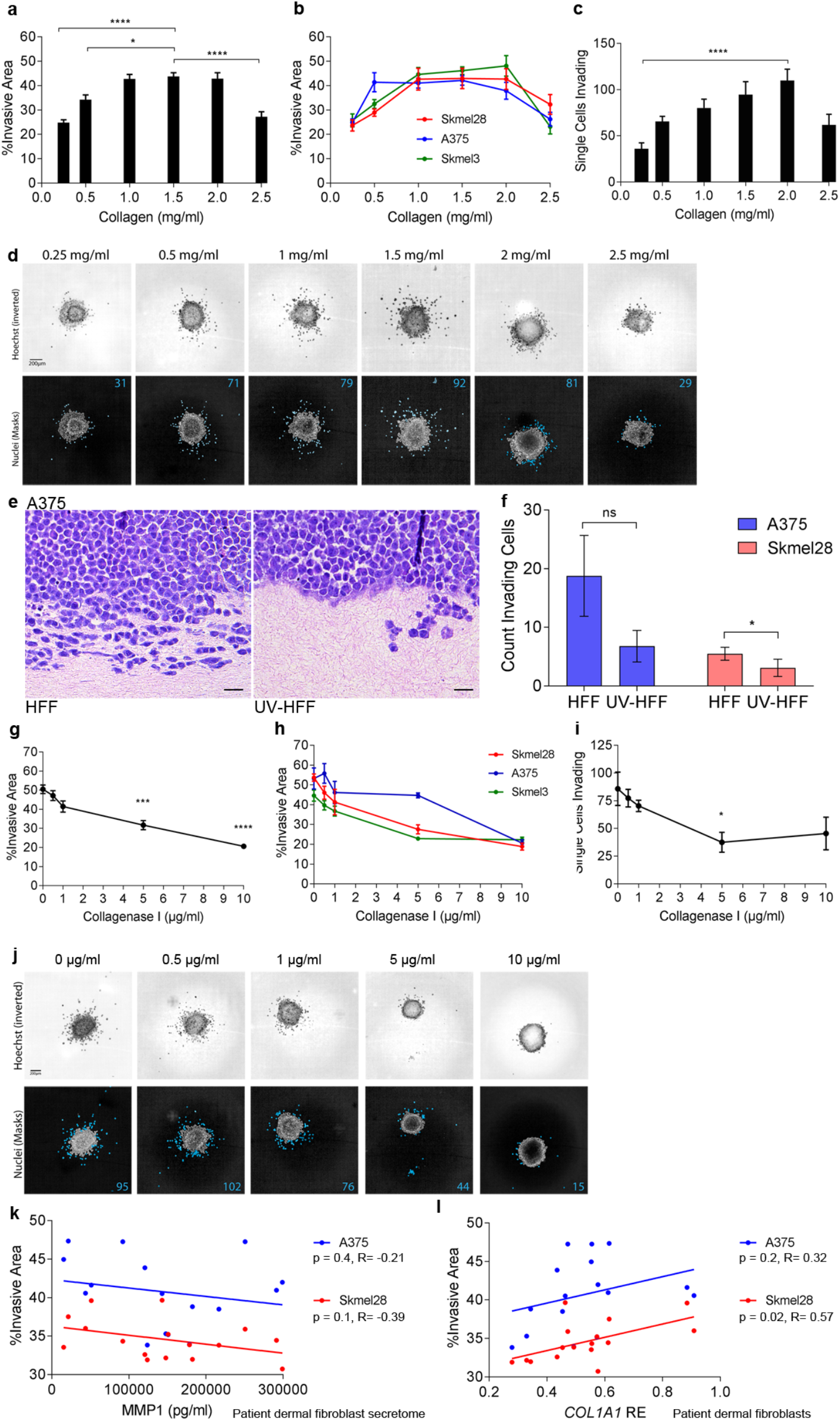
Low collagen quantity and integrity decrease melanoma cell invasion. **(a)** Mean and individual **(b)** melanoma spheroid invasion (Kruskal-Wallis with Dunn’s multiple comparison tests *p<0.05, ****p<0.0001) and **(c)** single cell invasion (Kruskal-Wallis with Dunn’s multiple comparison tests ****p<0.0001) in different collagen concentrations. **(d)** Representative images of spheroid and melanoma single cell invasion, top: Hoechst (inverted), bottom: invading cells. **(e)** A375 melanoma invasion H&E and **(f)** single cell invasion in organotypic dermal collagen HFF and UV-HFF constructs (Mann Whitney U *p<0.05, ns: not significant). **(g)** Mean and individual **(h)** melanoma invasion (Kruskal-Wallis with Dunn’s multiple comparison tests ***p<0.001,****p<0.0001) and **(i)** melanoma single cell invasion (Kruskal-Wallis with Dunn’s multiple comparison tests *p<0.05) by collagenase I concentration. **(j)** Representative images of spheroid and single cell invasion, top: Hoechst (inverted), bottom: invading cells. **(k)** Melanoma spheroid invasion by adult patient fibroblast secretome, MMP1 levels (Pearson correlation, red: Sk-mel-28 R =−0.39 p =0.1, blue: A375 R =−0.21 p =0.4) and **(l)** by *COL1A1* relative expression (RE) in fibroblasts (Pearson correlation, red: Sk-mel-28 R =0.57 p =0.02, blue: A375 R =0.32 p =0.2). Error bars: standard error of the mean (bar).

Since UVR compromises collagen integrity indirectly by damaging fibroblasts (Fig. 1), we exposed melanoma spheroids, embedded in equal concentrations of collagen matrices, to increasing concentrations of the enzyme collagenase I to mimic the effects of UVR exposure (Extended Data Fig. 2h). We found that melanoma invasion and single cell invasion significantly decreased in matrices exposed to higher doses of collagenase I (5μg/ml, p=0.0005, 10μg/ml, p<0.0001) (Fig. 2g-j, Extended Data Fig. 2i). These data show that collagen quantity and degraded collagen limit melanoma invasion.

To explore if adult fibroblasts regulate collagen degradation and melanoma invasion, we harvested adult dermal fibroblasts from different anatomic sites from tumour-adjacent normal skin of 8 patients and established cells lines (Extended Data Table 4, age median 69, range 34-77). We confirmed the patient dermal fibroblasts express and secrete varying levels of MMP1 (Extended Data Fig. 2j, k), and embedded melanoma spheroids in matrices of collagen mixed with the donor fibroblast secretome. We found melanoma spheroids were less invasive in matrices containing patient fibroblast secretomes with higher amounts of MMP1 (Methods, Fig. 2k). Importantly, *COL1A1* expression correlated strongly with melanoma invasion, (Sk-mel-28 p=0.02, R = 0.32, Fig. 2l). Finally, we confirmed that human fibroblasts, and not melanoma cells, are the main source of *MMP1*; and the melanoma cell lines do not express *COL1A1* or *COL1A2*^39^ (Extended Data Fig. 2l). Taken together, these data demonstrate human adult fibroblasts modulate collagen biology and melanoma invasion.

### Inhibition of MMP1 restores melanoma invasion

We studied if adult human fibroblasts from distinct anatomic sites affect the ECM and melanoma invasion differentially, and established fibroblast lines from chronic sun damaged (CSD) and sun-protected (noCSD) tumour-adjacent skin^34^ of two patients (age CSD = 77, age noCSD = 46). Donor fibroblasts were exposed to low doses of UVB (8x 100J/m^2^) to generate isogenic pairs of noCSD and noCSD-UV fibroblasts, CSD and CSD-UV fibroblasts and allowed 14 days recovery (Extended Data Fig. 2m). We confirmed noCSD-UV fibroblasts robustly increased MMP1 expression (fold change =2.03, p =0.002) and secretion (median MMP1: noCSD =6858 pg/ml, noCSD-UV =13292 pg/ml, fold change =1.98, p =0.03) compared to noCSD donor fibroblasts (Fig. 3a, b); and CSD-UV fibroblasts weakly upregulated *MMP1* expression (fold change =1.46, p =0.39), and secretion (median MMP1: CSD =9066 pg/ml, CSD-UV =13051 pg/ml, fold change =1.49, p =0.03). Importantly, noCSD-UV and CSD-UV fibroblasts did not increase *COL1A1* expression (Extended Data Fig. 2n). We then compared the effect of the fibroblast secretomes on melanoma spheroid invasion in the presence or absence of the MMP inhibitor Batimastat, which directly blocks the activity of MMPs. Consistent with a higher MMP1 expression, the noCSD-UV secretome significantly decreased melanoma invasion compared to the noCSD secretome (p=0.001), and importantly, Batimastat restored melanoma cell invasion (p=0.01) (Fig. 3c). Intriguingly, UVR damage or Batimastat treatment of the CSD fibroblast model (CSD-UV) only slightly modulated melanoma invasion (Fig. 3d), possibly indicating the effect of UVR damage to fibroblasts and *MMP1* expression is capped, and higher doses of MMP inhibition are required in highly expressing *MMP1*, CSD-UV fibroblasts.

**Figure 3.**
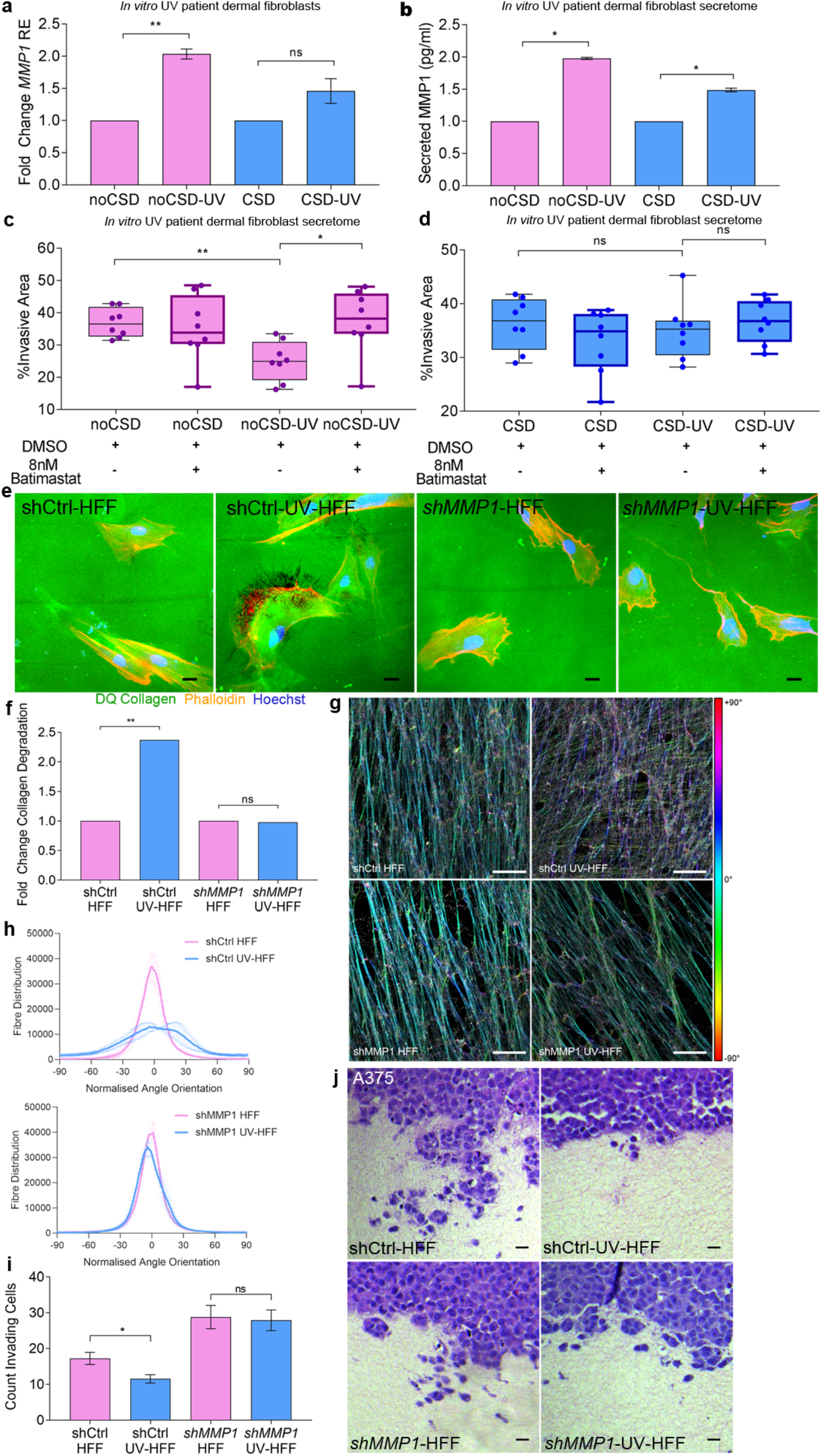
Inhibition of MMP1 restores melanoma invasion. **(a)** Fold change in *MMP1* relative expression (RE) in chronically UVR treated adult patient fibroblasts compared to untreated isogenic cell lines, noCSD: no chronic sun damage, pink, CSD: chronic sun damage, blue (Mann Whitney U **p<0.01, ns: not significant). **(b)** Fold change in secreted MMP1 in chronically UV treated adult fibroblasts compared to untreated isogenic cell lines, noCSD: no chronic sun damage, pink, CSD: chronic sun damage, blue, (Mann Whitney U *p<0.05). **(c)** Melanoma spheroid invasion in noCSD and isogenic noCSD-UV fibroblast secretomes in the presence of Batimastat or DMSO vehicle, (Mann Whitney U **p<0.01, *p<0.05). **(d)** Melanoma spheroid invasion in CSD and isogenic CSD-UV fibroblast secretomes in the presence of 8nM Batimastat or DMSO vehicle, ns: not significant. **(e)** Representative images of collagen degradation in shCtrl-HFF, shCtrl-UV-HFF, shMMP1-HFF and shMMP1-UV-HFF fibroblasts. Green: intact DQ collagen; red: phalloidin; blue: Hoechst. Size bars: 20 um **(f)** Fold change in collagen degradation of shCtrl-HFF and shMMP1-HFF (pink) and their isogenic chronic UVR cell lines shCtrl-UV-HFF and shMMP1-UV-HFF (blue), (Mann Whitney U **p<0.01, ns: not significant). **(g)** Immunofluorescence of fibronectin fibres in decellularised shCtrl-HFF, shCtrl-UV-HFF, shMMP1-HFF and shMMP1-UV-HFF derived ECM, colour coded for orientation of fibre, cyan represents mode, scale bar: 25 μm. **(h)** Quantification of fibre alignment distribution in shCtrl-HFF, shCtrl-UV-HFF, shMMP1-HFF and shMMP1-UV-HFF derived ECM. **(i)** Quantification of invading melanoma cells into organotypic dermal collagen constructs made with shCtrl-HFF and shMMP1-HFF (pink) and isogenic chronic UVR cell lines shCtrl-UV-HFF and shMMP1-UV-HFF (blue), (Mann Whitney U *p<0.05, ns: not significant). **(j)** Representative images of A375 invasion into organotypic constructs stained with H&E. Error bars represent standard error of the mean (bar).

Since MMP1 specifically cleaves COL1A1^28^, we generated isogenic shCtrl-HFF, shCtrl-UV-HFF, *shMMP1*-HFF and *shMMP1*-UV-HFF lines from human foreskin fibroblasts to compare collagen degradation in the absence of acute UVR exposure (Extended Data Fig. 3b, c). In keeping with a higher expression of MMP1, shCtrl-UV-HFF fibroblasts degraded more collagen, while *shMMP1*-UV-HFF did not increase collagen degradation compared to *shMMP1*-HFF fibroblasts (Fig. 3e, f). The specific role of MMP1 was validated with an additional knockdown with an siRNA targeting MMP1 (Extended Data Fig. 3d, e, f). Additionally, we found shRNA targeting shMMP2 did not modify collagen degradation (Extended Data Fig. 3g, h); and knockdown of MMP1 restored the alignment of fibres in UV-HFF matrices. (Fig. 3g, h, Extended Data Fig. 3i). Furthermore, organotypic invasion assays with matrices generated with shCtrl-HFF, shCtrl-UV-HFF, sh*MMP1*-HFF or sh*MMP1*-UV-HFF fibroblasts, showed melanoma invasion was decreased in the shCtrl-UV-HFF constructs (p=0.03), and not in sh*MMP1*-HFF or sh*MMP1*-UV-HFF fibroblast matrices (Fig. 3i, j). Knockout of MMP1 restored collagen and fibronectin levels in UV-HFF constructs to similar levels as HFF (Extended Data Fig. 3j, k, l). Altogether, these data demonstrate fibroblast-secreted MMP1 degrades collagen, limiting melanoma invasion.

### Collagen degradation decreases primary melanoma invasion and improves survival

If the amount and integrity of collagen restrict single cell invasion, patients with primary cutaneous melanoma invading in a less collagenous dermis should live longer than patients with more dermal collagen. Compared to young fibroblasts and dermis, aged and UVR-protected fibroblasts in an aged ECM drive melanoma invasion and metastasis^40,41^, so we restricted our study to three international cohorts of older primary cutaneous melanoma patients. We determined the proportion of melanoma cells invading in the ECM at the invasive front (IF), the amount of collagen and the degree of ECM degradation (solar elastosis) in tumour-adjacent dermis (Fig. 4a, Extended Data Fig. 4a). We found patient samples with more solar elastosis (less collagen in tumour-adjacent skin), had fewer invading cells at the IF (Fisher exact test, p =2.25×10^−5^ Fig. 4b, Extended Data Fig. 4b). Critically, melanoma specific survival (MSS) was significantly improved in patients with less invasion in multivariate analyses (Fig. 4c, Extended Data Fig. 4c, d, Extended Data Table 6). Intriguingly, solar elastosis was not as powerfully associated with better outcome (p =0.9, Fig. 4d, Extended Data Fig. 4e, f, Extended Data Table 6). To explain this difference, we hypothesised that collagen at the IF, rather than collagen degradation in the tumour-adjacent dermis, would be a better biomarker of survival. Further analysis confirmed that collagen at the IF strongly correlated to single cell invasion (Spearman R 0.5, p<0.0001, Fig. 4e), MSS and progression free survival (PFS; Fig. 4f, Extended Fig. 4g, Extended Data Table 6). Furthermore, consistent with invasion data in Fig 2j, TCGA primary cutaneous melanomas expressing low *COL1A1* showed improved survival (Fig. 4g, Extended Data Table 6). These data suggest that primary melanomas invading in collagen-poor matrices require new collagen synthesis in order to invade successfully. To confirm this, we studied collagen at the IF specifically, in short-term and long-term survivors. We confirmed melanomas arising at chronic sun damaged sites (CSD) with less collagen in the tumour-adjacent dermis and shorter survival (MSS <5 years), increased collagen deposition at the IF and tumour cell invasion (Fig. 4h, 4i, Extended Data Table 6).

**Figure 4.**
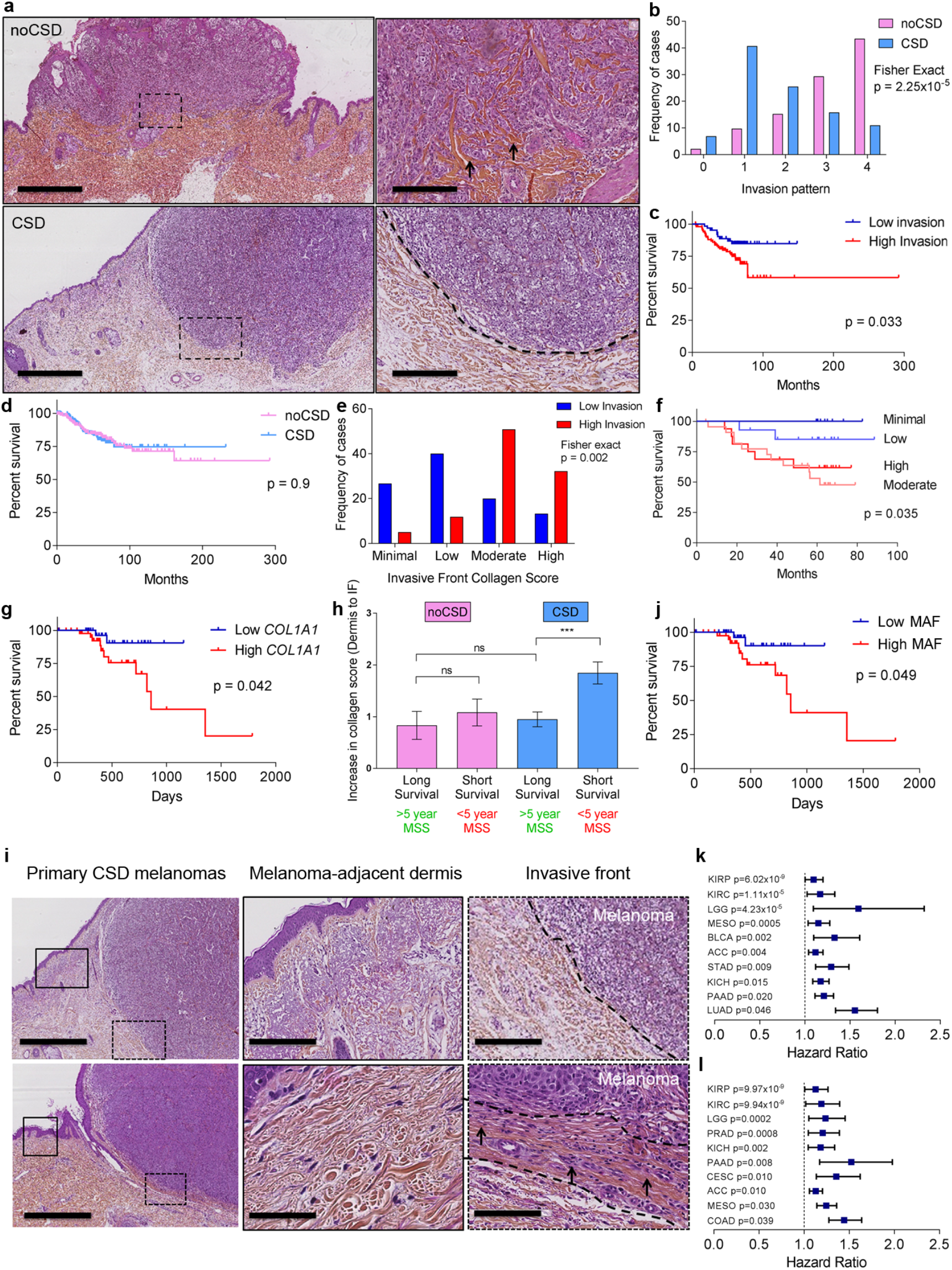
Low Collagen correlates with low invasion and improved outcome in aged melanoma patients. **(a)** H&E top panel: Primary cutaneous melanoma and inset (box) with single cell invasion (arrows) in a collagen rich dermis, collagen: red (scale bars left: 4000μm, right: 400μm. Lower panel: Melanoma in a collagen-poor dermis and inset (box), no single cell invasion; collagen: red (scale bars left: 3000μm, right: 300μm). **(b)** Histogram displaying melanoma invasion at the invasive front (IF) in noCSD and CSD melanomas (B and C cohorts, n=170). **(c)** Kaplan-Meier of melanoma specific survival (MSS) in prominent (high, red) and minimal (low, blue) melanoma invasion at the IF (Log Rank test, B and C cohorts, n= 168). **(d)** Kaplan-Meier of MSS in melanoma invading in CSD (blue) and noCSD (pink) dermis. (Log Rank test, B and C cohorts, n=331) **(e)** Histogram displaying collagen quantity at the IF in highly invasive (red) and minimally invasive (blue) melanoma (C cohort, n=89). **(f)** Kaplan-Meier of MSS by collagen quantity at the IF (Log Rank test, C cohort, n=62). **(g)** Kaplan-Meier of MSS by *COL1A1* expression in aged (>54) primary cutaneous melanoma (Log Rank test, TCGA cohort, n=80). **(h)** Fold increase in collagen deposition at the IF of noCSD and CSD melanomas by MSS (Mann Whitney U ***p<0.001 C cohort n=90). **(i)** H&E top panel: left CSD melanoma, middle: from box inset: tumour-adjacent dermis; right: from dashed box inset: IF (dashed line, scale bars: 2000μm, 300μm, 300μm). Bottom: left CSD melanoma, middle: from box inset: tumour-adjacent dermis; right: from dashed line box inset: IF between dashed lines, arrows: melanoma invasion, (scale bars: 2000μm, 70μm, 200μm). **(j)** Kaplan-Meier of MSS by MAF signature score in aged (>54) primary cutaneous melanoma cohort (Log Rank test, TCGA cohort, n=80). **(k)** Hazard ratio and 95% CI for OS, and PFS **(l)** in univariate Cox regression of *COL1A1* expression by cancer type in PANCAN TCGA (ACC, Adrenocortical carcinoma, BLCA, Bladder urothelial carcinoma, CESC, Cervical and endocervical cancers, COAD, Colon adenocarcinoma, KICH, Kidney chromophobe, KIRC, Kidney renal clear cell carcinoma, KIRP, Kidney renal papillary cell carcinoma, LGG, Brain lower grade glioma, LUAD, Lung adenocarcinoma, MESO, Mesothelioma, PAAD, Pancreatic adenocarcinoma PRAD, Prostate adenocarcinoma, STAD, Stomach Adenocarcinoma). Risk tables for all Kaplan-Meier analyses in Extended Data Table 6.

To further explore the association between collagen and survival, we investigated if melanoma-associated fibroblasts (MAFs) in primary melanomas increase collagen to sustain invasion. For this, we extracted the gene expression signature from single cell RNAseq^39^ and confirmed *COL1A* is expressed by MAFs (Extended Data Fig. 4h). We then tested if increased expression of MAFs in primary melanomas correlates with outcome and demonstrate a higher expression of MAF genes is associated with poor survival (Fig. 4j). Furthermore, we show it is the expression of collagen genes specifically within the MAF expression signature that impacts survival (Extended Data Fig. 4i, Extended Data Table 6).

Solid cancers synthesise high amounts of ECM proteins and COL1A1; and ECM remodelling promotes primary tumour progression and metastasis^42,43^. Therefore, we designed a tumour-agnostic approach to test the potential of *COL1A1* expression as a biomarker for primary pan-cancer survival, which revealed young and aged patients with primary cancers expressing high levels of *COL1A1* are at greater risk of death and have shorter progression free survival (PFS; Figure 4k, l, Extended Data Fig. 4j, k).

## Discussion

Multiple *in vivo* studies confirm UVR cooperates with oncogenic mutations to increase the incidence and penetrance of disease^10,44^. However, whether UVR exposure affects the odds of survival has not been comprehensively investigated and there are contradictory studies finding sun exposure inferred by anatomic site^13^, history of sunburn^14,18^ or the presence of UVR-induced dermal degradation^11,15^ can affect outcome. The majority of melanoma deaths affect the elderly^7^, and age is strongly associated with accumulated sun exposure^12^. We investigated if pre-existing UVR damage affects melanoma survival and found low collagen quantity and integrity limit melanoma invasion. UVR damage to fibroblasts degrades collagen and the ECM, delaying melanoma progression. We confirmed our *in vitro* results, showing that in aged primary cutaneous melanomas, single tumour cells invading the dermis and collagen at the IF robustly predict poor survival. Paradoxically, this study finds UVR damage to the dermis destroys collagen, limiting invasion and improving outcome, unless tumours increase the production of collagen at the IF, providing the structural support for melanoma invasion. Together with recent work showing UVR-protected aged fibroblasts^40^ and ECM^41^ drive melanoma metastasis, this study strongly implicates the physical composition and structure of the aged tumour microenvironment as key to primary melanoma progression. We therefore infer from these joint studies that excessive old age mortality particularly affects patients with tumours arising at anatomic sites with preserved dermal collagen, or sun-protected skin. In contrast, UVR damage modifies the dermis and decreases collagen content as we age. As melanomas arising in collagen-poor, sun-damaged skin require collagen to invade, we show that collagen deposition at the invasive edge of the tumour is an independent, robust biomarker of survival. Conveniently, the deposition of collagen at the IF can be scored simply from haematoxylin and eosin stains, making this an ideal biomarker.

Melanomas with more UVR damage accumulate more mutations and neoantigens, possibly eliciting stronger immune responses^45,46^. However, we show a proportion of UVR melanomas have a better prognosis due to collagen degradation, independently of the mutation burden, tumour and stromal cell immunogenicity, which should be considered when evaluating responses to adjuvant immunotherapy. One possibility is to prioritise adjuvant care according to single cell invasion, collagen at the IF and risk of death. Supporting this rationale, recent evidence shows collagen density modifies the immune milieu of breast cancers, limiting T cell responses^47^.

The accumulation of a collagenous ECM, increasing stiffness, leads to poor prognosis and lack of response to therapies in other cancers^42^. Alterations of the ECM dynamics are a hallmark of cancer, able to deregulate cancer and stromal cells, promote cell transformation and the pro-metastatic niche, which becomes rich in vasculature and tumour-promoting inflammation^48^. Clinical trials with drugs inhibiting MMPs and limiting ECM remodeling have yielded negative and sometimes deleterious results. One possible explanation for this failure, based on our study in melanoma, is that inhibition of collagen degradation may support tumour invasion. Collagen can drive cancer cell de-differentiation^22,49–51^ and is often found in areas of active epithelial cancer invasion, facilitating migration^50,52^.

Ageing is associated with less collagen deposition, more degradation and higher overall cancer incidence and mortality in multiple tissues, including skin. One possibility is the collagen decrease in aged tissue, which could lead to less aggressive cancer, is offset by a decrease in ECM structural fitness, collagen and matrix organisation, and pro-tumourigenic signalling with age^43^. This work suggests new collagen synthesis by the TME is a critical regulator of aged primary melanoma progression, a feature that could drive poor outcome in other aged cancers. Critically, our data shows collagen expression is associated in multiple solid epithelial and non-epithelial primary tumours with shorter PFS in all ages, possibly due to direct modulation of invasion.

## Extended Data Supplementary Figures

**Extended Data Figure 1.**
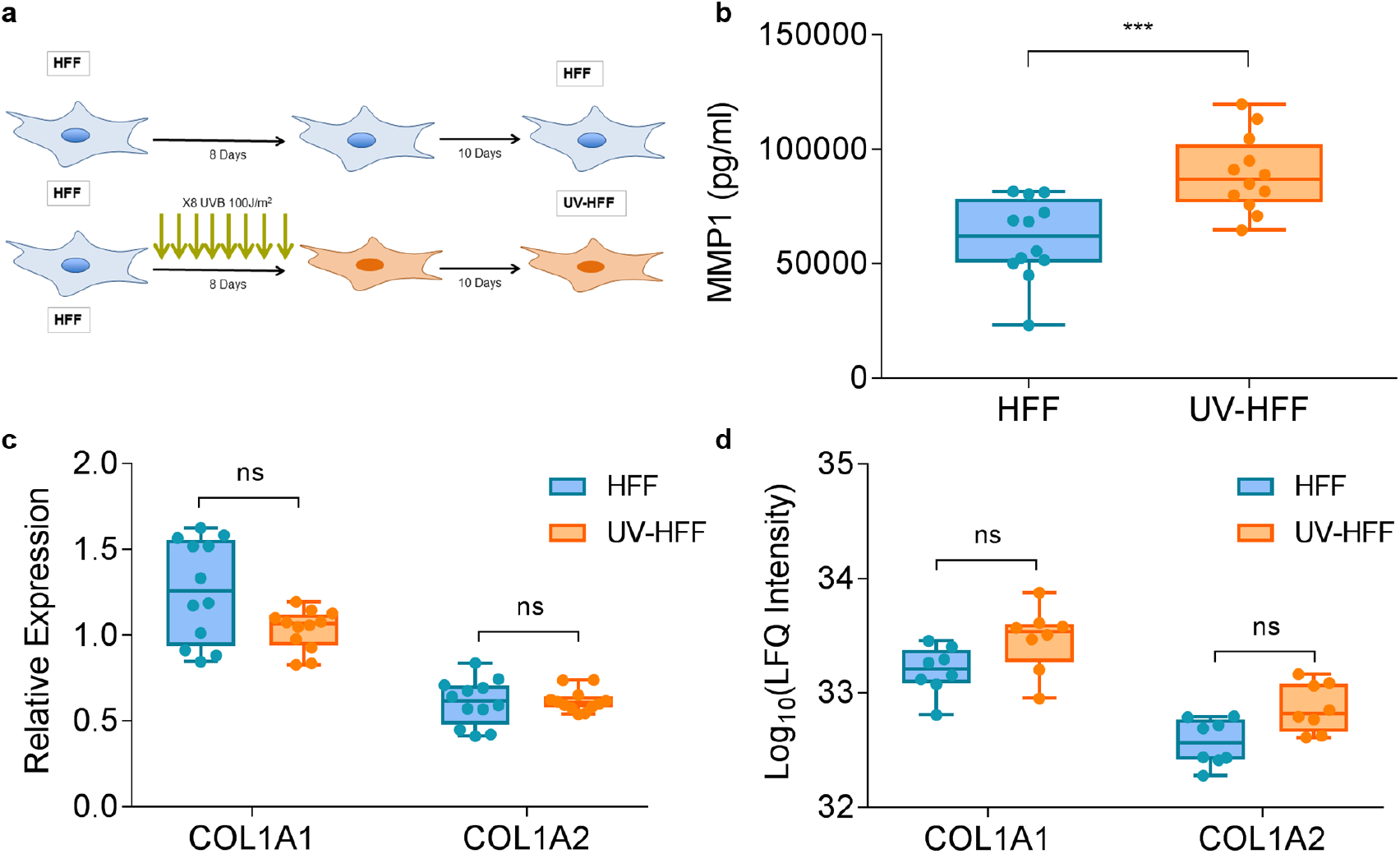
UVR damaged fibroblasts upregulate MMP1 but not COL1A1. **(a)** Graphic representation of the generation of isogenic HFF and UV-HFF by chronic UVR treatment. **(b)** Quantification of MMP1 in the secretome of HFF and UV-HFF fibroblasts, (Mann Whitney U ***p<0.001). **(c)** Relative Expression of *COL1A1* and *COL1A2* in HFF and UV-HFF fibroblasts, (Mann Whitney U ns: not significant). **(d)** Label free quantification (LFQ) of COL1A1 AND COL1A2 in the HFF and UV-HFF matrisome by mass spectrometry, (Mann Whitney U ns: not significant). All data represents duplicate samples collected from biological replicate cell lines, box plot showing all replicate points. Error bars: standard error of the mean (bar).

**Extended Data Figure 2.**
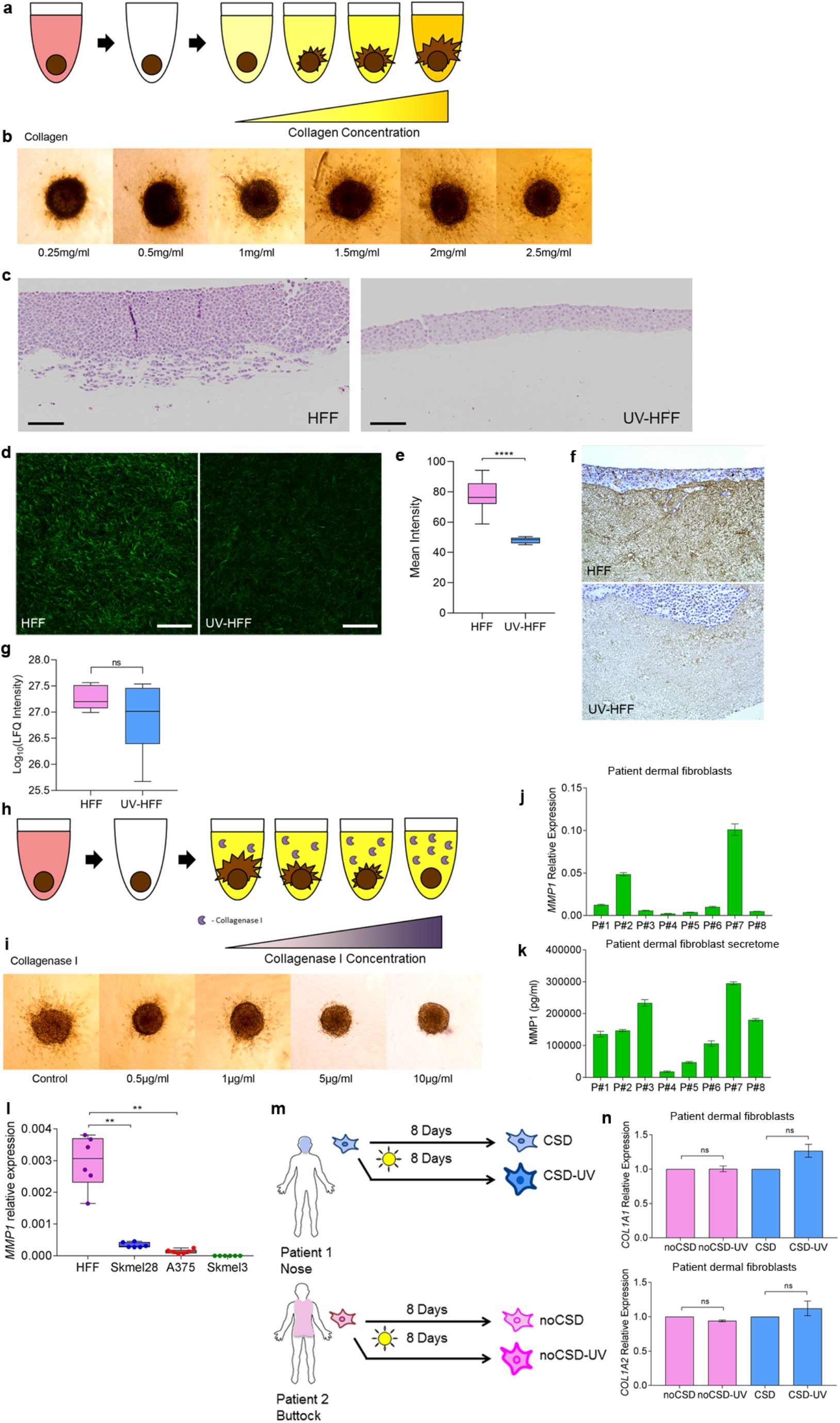
Melanoma invasion decreases in low amounts and degraded collagen. **(a)** Graphical representation of spheroid model of collagen concentration gradient. **(b)** Representative Sk-mel-28 spheroid invasion in collagen gradient. **(c)** Representative H&E photomicrographs of UV and UV-HFF constructs with melanoma cells (scale bar: 150 μm) **(d)** Second harmonic generation (SHG) imaging of collagen fibres in organotypic dermal collagen HFF and UV-HFF constructs, scale bar: 50 μm. **(e)** Quantification of collagen from SGH images in HFF and UV-HFF, (Mann Whitney U ****p<0.0001). **(f)** Fibronectin IHC staining in organotypic dermal collagen HFF and UV-HFF constructs. **(g)** Label free quantification (LFQ) of elastin (ELN) in the HFF and UV-HFF matrisome by mass spectrometry (Mann Whitney U ns: not significant). **(h)** Graphical representation of spheroid model of collagenase I concentration gradient. **(i)** Representative Sk-mel-28 spheroid invasion in collagenase I gradient. **(j)** Relative expression (RE) of *MMP1* in panel of eight healthy adult dermal fibroblasts cell lines. **(k)** Quantification of MMP1 in the secretome of eight adult fibroblast cell lines. **(l)** Relative expression of *MMP1* in melanoma cell lines (Sk-mel-28, A375, Sk-mel-3) compared to HFF fibroblasts, (Mann Whitney U **p<0.01). **(m)** Graphical representation of isogenic chronic UVR model in two adult fibroblast lines. **(n)** Fold change in expression of *COL1A1* and *COL1A2* in chronic UVR fibroblasts normalised to isogenic untreated cell lines, (Mann Whitney U ns: not significant). Error bars: standard error of the mean (bar).

**Extended Data Figure 3.**
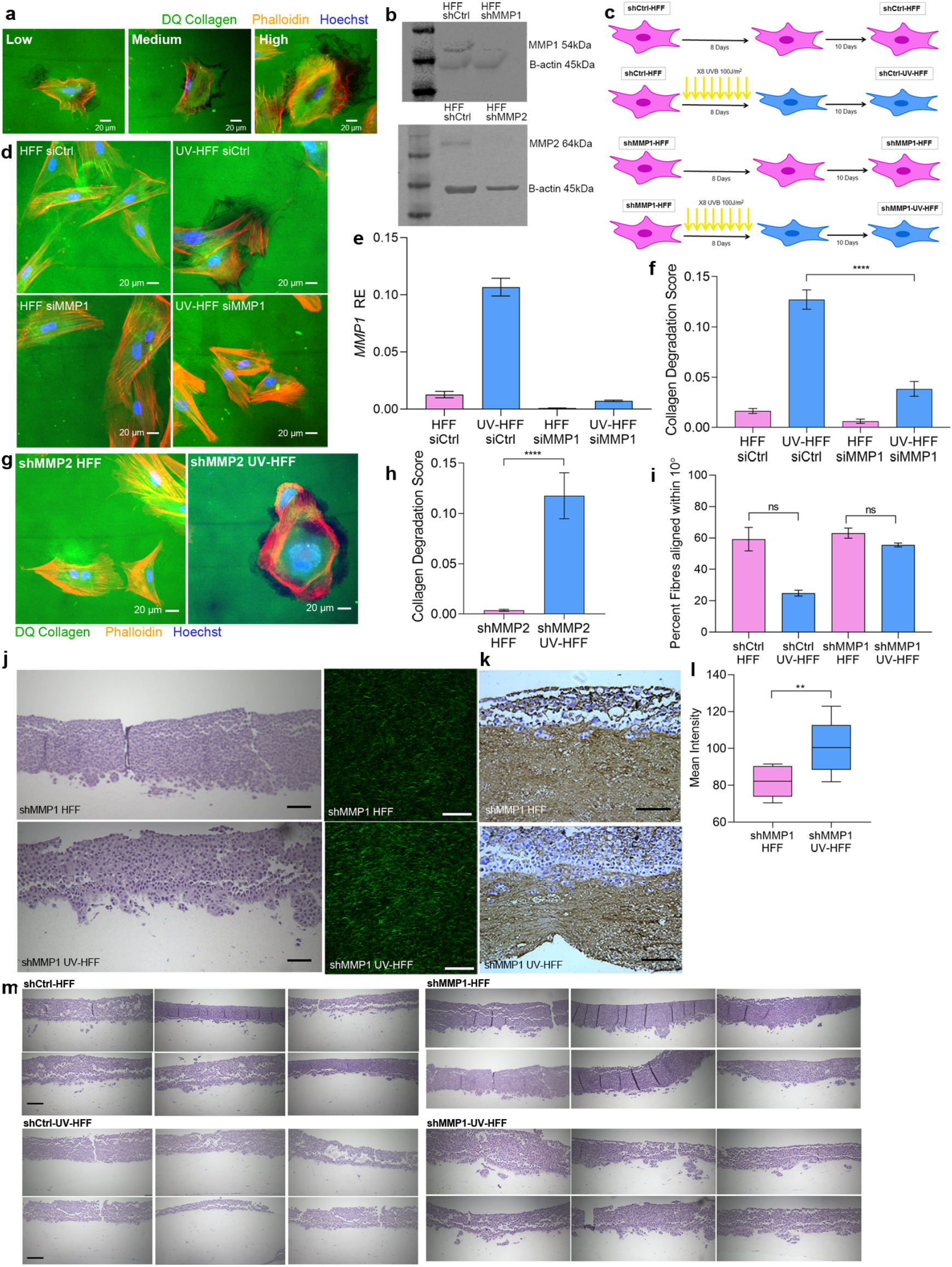
UVR damage to human dermal fibroblasts upregulate MMP1 but not COL1A1. **(a)** Representative images of collagen degradation scores as outlined in methods. **(b)** Western blots validating knockdown of MMP1 and MMP2 in shRNA cell lines. **(c)** Graphical representation of isogenic chronic UVR model in shCtrl and shMMP1 HFF fibroblasts. **(d)** Representative images of collagen degradation in siCtrl-HFF, siCtrl-UV-HFF, siMMP1-HFF and siMMP1-UV-HFF fibroblasts. Green: intact DQ collagen; red: phalloidin; blue: Hoechst. Size bars: 20 μm. **(e)** Validation of siRNA effect on *MMP1* relative expression (RE) by qPCR in collagen degradation assay. **(f)** Quantification of collagen degradation of siCtrl-HFF and siMMP1-HFF (pink) and their isogenic chronic UVR cell lines siCtrl-UV-HFF and siMMP1-UV-HFF (blue), (Mann Whitney U ****p<0.0001). **(g)** Representative images of collagen degradation in shMMP2-HFF, and shMMP2-UV-HFF fibroblasts. Green: intact DQ collagen; red: phalloidin; blue: Hoechst. Scale bars: 20 μm. **(h)** Quantification of collagen degradation of shMMP2-HFF and their isogenic chronic UVR cell line shMMP2-UV-HFF (blue), (Mann Whitney U ****p<0.0001). **(i)** Quantification of fibres within 10° of mode orientation in shCtrl-HFF, shCtrl-UV-HFF, shMMP1-HFF and shMMP1-UV-HFF derived ECM by fibronectin immunofluorescence (shCtrl HFF vs shCtrl UV-HFF p=0.1 (n=3), ns: not significant) **(j)** Representative H&E images of shMMP1-HFF and shMMP1-UV-HFF fibroblasts, scale bars: 75 μm (left) and second harmonic generation (SHG) imaging of collagen fibres in organotypic dermal collagen shMMP1-HFF and shMMP1-UV-HFF constructs (right), scale bars: 50 μm. **(k)** Fibronectin IHC staining in organotypic dermal collagen shMMP1-HFF and shMMP1-UV-HFF constructs, scale bars: 50 μm. **(l)** Quantification of collagen from SGH images in shMMP1-HFF and shMMP1-UV-HFF (Mann Whitney U, **p<0.01). **(m)** Representatives H&E photomicrographs of shCtrl-HFF (four top left images), shCtrl-UV-HFF (four top right images), shMMP1-HFF (four bottom left images) and shMMP1-UV-HFF (four bottom right images) derived ECM constructs with invading melanoma cells, scale bars: 100 μm. Error bars: standard error of the mean (bar).

**Extended Data Figure 4.**
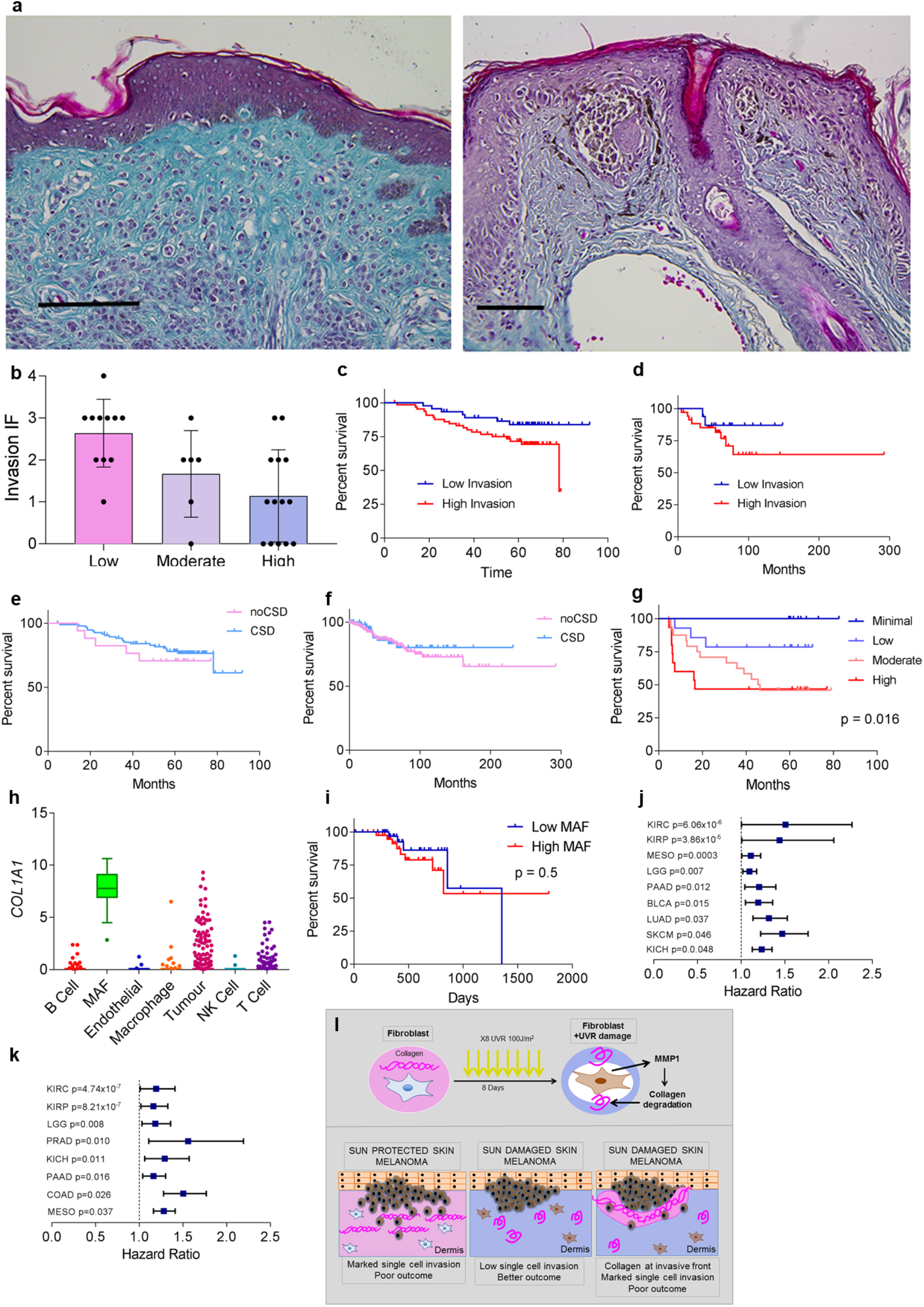
Collagen at the invasive front or primary invasive cutaneous melanoma drives melanoma invasion and poor outcome. **(a)** Trichrome of Masson stain of collagen in sun protected skin (left, scale bar 200 μm) and sun damaged skin (right, scale bar 200 μm). **(b)** Correlation between ECM degradation of the adjacent dermis and invasive score of melanoma tumour cells in the A cohort. **(c)** Kaplan-Meier of melanoma specific survival (MSS) in prominent (high, red) and minimal (low, blue) melanoma invasion at the IF (C cohort, n=112). **(d)** Kaplan-Meier of melanoma specific survival (MSS) in prominent (high, red) and minimal (low, blue) melanoma invasion at the IF (B cohort, n=51). **(e)** Kaplan-Meier of MSS in melanoma invading in CSD (blue) and noCSD (pink) dermis (C cohort, n=113). **(f)** Kaplan-Meier of MSS in melanoma invading in CSD (blue) and noCSD (pink) dermis (B cohort, n=216). **(g)** Kaplan-Meier of progression free survival by collagen quantity at the IF (Log Rank test, C cohort, n=63). **(h)***COL1A1* expression by cell type in metastatic melanoma single cell RNA-seq (Tirosh *et al.* Science 2016). **(i)** Kaplan-Meier of MSS by MAF signature score without collagen genes in aged (>54) primary cutaneous melanoma cohort (Log Rank test, TCGA cohort, n=80). **(j)** Hazard ratio and 95% CI for OS, and PFS **(k)** Univariate Cox regression of *COL1A1* expression by cancer type in PANCAN TCGA in aged population (≥55 years) (BLCA, Bladder urothelial carcinoma, COAD, Colon adenocarcinoma, KICH, Kidney chromophobe, KIRC, Kidney renal clear cell carcinoma, KIRP, Kidney renal papillary cell carcinoma, LGG, Brain lower grade glioma, LUAD, Lung adenocarcinoma, MESO, Mesothelioma, PAAD, Pancreatic adenocarcinoma PRAD, Prostate adenocarcinoma, SKCM, Skin Cutaneous Melanoma). **(l)** Graphic summary of study. Risk tables for all Kaplan-Meier analyses in Extended Data Table 6. Error bars: standard error of the mean (bar).

## Extended Data Supplementary Tables

**Supplementary Table 1 –** Differentially expressed genes by COSMIC signature 7 mutation count in healthy adult dermal fibroblasts

**Supplementary Table 2 –** Top 10 Reactome pathways significantly enriched by genes differentially expressed by signature 7 mutation count

**Supplementary Table 3 –** Genes from Reactome Extracellular Matrix Organisation pathway differentially expressed by Signature 7 mutations

**Supplementary Table 4 –** Genes from Reactome Extracellular Matrix Organisation pathway differentially expressed

**Supplementary Table 5 –** Clinical cohort details

**Supplementary Table 6 –** Cox regression of clinical, invasion and collagen variables, risk tables for Kaplan-Meiers

## METHODS

### Cell lines and patient fibroblasts

Human foreskin fibroblasts (HFF) were purchased from ATCC (ATCC SCRC-1041). Three melanoma cell lines, Sk-mel-28, Sk-mel-3 and A375 were obtained from ATCC. All cell lines were cultured in DMEM (Gibco) supplemented with 10% FCS (Sigma Aldrich), 1X Glutamax (Gibco), 100U/mL Penicillin and Streptomycin (Gibco), 1mM Sodium pyruvate (Gibco). Cells lines were cultured at 37°C in 5% CO_2_ with medium replaced as required. Cell lines were tested every fortnight for Mycoplasma using LookOut Mycoplasma PCR Detection Kit (Sigma Aldrich). Cell line identity was confirmed using STR profiling.

### Patient fibroblasts

A prospective cohort of patient fibroblast cultures was established from redundant skin acquired during surgical resection of the wide local excision of healthy skin from melanoma patients treated at the tertiary referral cancer hospital in Cohort A, in vitro studies. Ethical approval was granted by the the local Biobank committee (reference number will be included). We obtained informed consent from all participants. The hypodermis of whole skin samples was removed scraping with a scalpel, and the residual specimen was incubated overnight in Dispase (Gibco) at 4°C to separate the epidermis and dermis. The dermis was digested in Collagenase I (Gibco) in DMEM (without FCS) (Gibco) at 37°C for 6 hours and then filtered through 70μm filter to remove the residual debris. Dermal cells were spun at 300xg and re-suspended in DMEM 20% FCS, cultured in 20% FCS DMEM until they became confluent, and stained for vimentin (Abcam). The level of solar elastosis of the redundant skin collected was scored using previously well-established methods^1^.

### Whole exome sequencing of patient fibroblasts

Whole exome sequencing of fibroblasts was performed by Novogene (Novogene (UK) Company Ltd.). Exome capture was performed with the SureSelect Human All Exon v6 kit (Agilent) and sequenced on the Illumina HiSeq platform. Sequencing reads were trimmed using Trimmomatic^2^, aligned to the hg38 reference genome using BWA^3^ and duplicate reads were marked using Picard Tools (http://broadinstitute.github.io/picard). Somatic mutations were called using the Varscan 2 pipeline^4^. Identified somatic variants were annotated using Variant Effect Predictor^5^ and variants present in dbSNP (but not in the COSMIC database) were excluded.

### Lentiviral shRNA transfection

Knockdown of *MMP1* expression in HFF cells was performed using shRNA Lentiviral Particles (Santa Cruz Biotechnology) and siGENOME siRNA (Horizon Discovery). For *MMP1* shRNA knockdown MMP-1 shRNA (h) Lentiviral particles (sc-41552-V) were used alongside a scramble control, Control shRNA Lentiviral Particles A (sc-108080), and copGFP Control Lentiviral Particles (sc-108084) were used to measure transduction efficiency. 5×10^4^ cells were cultured in cell culture media with 5ug/ml Polybrene (Santa Cruz Biotechnology). Lentiviral particles were added to cells and incubated overnight. Media containing lentiviral particles and Polybrene was removed and incubated in DMEM overnight before preforming selection of transfected cells using increasing concentration of puromycin over 72 hours. Once cells were stably growing in puromycin, cells were cultured as normal in DMEM. For *MMP1* siRNA siGENOME Human MMP1 siRNA (D-005951-02-0002) and Non-Targeting siRNA #4 (D-001210-04-05) were used with DharmaFECT 1 Transfection Reagent according to manufactures protocols, with a final siRNA concentration of 25nM. Knockdown of MMP2 was performed with MMP-2 shRNA (h) Lentiviral particles (sc-29398-V) as above. Knockdown of all gene expression was validated by qPCR and western blot.

### UV treatment and CSD model

Cell lines were treated with UVB using a Bio-Sun UV irradiation system (Vilber Loumat). For chronic treatment the dose 100J/m^2^ was used as it represents a physiologically relevant dose of UVB that would penetrate the dermis between 1 to 5 minimal erythema dose (MED)^6^. To create isogenic *in vitro* chronic UV damaged UV-HFF and *shMMP1*-UV-HFF, noCSD-UV, CSD-UV cell lines, 1×10^6^ HFF, *shMMP1*-HFF, or patient dermal fibroblasts were cultured in 100mm dishes in phenol free DMEM 1% FCS. All fibroblasts were treated every 24 hours with 100J/m^2^ UVB for 8 consecutive days. Following the UV treatments, the medium was changed to DMEM 10% FCS and cultured for one week. Isogenic untreated control cell lines (HFF, *shMMP1*-HFF, noCSD, CSD) were cultured in identical conditions without the UVB treatments. Each HFF condition was created in biological duplicates.

### Secretome collection

To collect secretomes 1×10^6^ cells were plated in a 100mm dish and cultured for 72 hours in DMEM without FBS to limit cell proliferation. Secretomes were collected in duplicate, aliquoted and stored at −80°C until used.

### Batimastat treatment

MMP inhibitor Batimastat (Sigma Aldrich) was resuspended in DMSO at 15mg/mL and a stock was diluted to 1mM. Batimastat was added to fibroblast secretomes to a final concentration of 8nM in secretome volume and control secretomes had equal volume of DMSO added as a vehicle control.

### RNA-sequencing data analysis

RNA-seq data from ENA project PRJEB13731 (https://www.ebi.ac.uk/ena/browser/view/PRJEB13731); also at https://www.ebi.ac.uk/arrayexpress/experiments/E-MTAB-4652/) were downloaded. Data are single-end RNA-seq from short term cultivated fibroblasts sequenced on an Illumina HiSeq 2000 sequencer. Two samples had been obtained from each individual from different locations (B=buttock, not UV exposed; S=shoulder, UV exposed). Sequencing reads were trimmed^2^ and aligned to the human reference genome (GRCh37) using STAR^7^. Production of analysis-ready reads was conducted according to the Broad Institute Best Practices pipeline (https://gatkforums.broadinstitute.org/gatk/discussion/3892/the-gatk-best-practices-for-variant-calling-on-rnaseq-in-full-detail) using GATK v3.2 (http://www.broadinstitute.org/gatk). Somatic single nucleotide variants (SNVs) in each patient matched sample in regions annotated as protein-coding only (based on Ensembl Homo_sapiens.GRCh37.87) were identified with the mutation calling algorithm MuTect v1 using the other sample as the comparator ^8^. Identified somatic variants were annotated using Variant Effect Predictor^5^ and common variants were excluded.

COSMIC mutational signatures v2 (https://cancer.sanger.ac.uk/cosmic/signatures_v2) were identified using the MutationalPatterns ^9^ package (version 1.8.0) in R (version 3.5.1, RStudio v1.2.5001, RStudio Inc). Differential expression analysis was performed using the DESeq2 package (version 1.22.2, ^10^) in R (version 3.5.1). Reads counts of genes were filtered for genes expressed in fibroblasts by removing any gene with less than 100 counts across all samples. For pathway enrichment analysis genes that were significantly differentially expressed in by signature 7 mutation count (FDR p-value < 0.1) were compared against the Reactome Database^11^ using the Molecular Signatures Database v7.0 (Msigdb, Broad Institute, https://www.gsea-msigdb.org/gsea/msigdb/index.jsp).

### Fibroblast extracellular matrix (ECM) production

ECM was constructed as previously described with minor modifications^12^. Cell culture dishes were coated with 0.2% sterile gelatin, fixed with 1% glutaraldehyde and quenched with 1M glycine in PBS (pH7). Fibroblasts were cultured on gelatin plates in normal 10% FCS DMEM containing 50 μg/ml ascorbic acid for 8 days. Cells were lysed with extraction buffer (20mM NH_4_OH, 0.5% Triton-X100 in PBS) and washed thoroughly with PBS containing calcium and magnesium. DNA was digested with 10μg/ml DNase I (Roche) and washed. For mass spectrometry the ECM was collected with a lysis buffer (100 mM TrisHCl pH 7.5, 4% SDS, 100 mM DTT) and collected with scraper, sonicated and boiled at 95°C, followed by centrifugation (16,000 g, 15 min) and collection of the supernatant, stored at −80°C until use.

### Immunofluorescence and ECM fibre analysis

For immunofluorescence and ECM fibre analysis, fibroblasts derived matrices from above, after DNase digestion matrices, were fixed in 4% paraformaldehyde, blocked, and stained for Fibronectin following the previously published protocol^13^. Primary antibody to Fibronectin (1:200, F3648, Sigma Aldrich) and secondary Goat anti-Rabbit Alexa Fluor 488 (1:2000, Thermo Fisher) were incubated for 1 hour at room temperature. The immunofluorescence imaging was performed using a Carl Zeiss LSM880 inverted confocal microscope with 63x NA 1.4 oil objectives lens controlled by ZEN black software. The images were acquired with 488nm illumination laser line from an Argon laser (Lasos) and the emission spectrum range from 500 to 550nm collected with a PMT detector (Zeiss). Z-series optical sections were collected with a step-size of 0.5 micron driven by Piezo stage (Zeiss). Fibre orientation analysis was performed using ImageJ OrientationJ plugin^14^ as previously described^13^. Maximal projection of three individual z-stacks for each condition were analysed.

### Mass spectrometry sample preparation and analysis

ECM protein lysates were separated on a 4–12% gradient NuPAGE Novex Bis-Tris gel (Life Technologies). Each sample was cut into 3 slices and in-gel digested with trypsin (Promega) as previously described^15^. Digested peptides were desalted by C18 StageTip^16^, acetonitrile was removed by speed vacuum, and peptides were resuspended in 1% trifluoroacetic acid, 0.2% formic acid. Peptides were injected into an EASY-nLC (Thermo Fisher Scientific) coupled online to an Orbitrap Fusion Lumos mass spectrometer (Thermo Fisher Scientific), separated using a 20 cm fused silica emitter (New Objective) packed in house with reversed-phase Reprosil Pur Basic 1.9 μm (Dr Maisch GmbH) and eluted with a flow of 300 nl/min from 5% to 30% of buffer (80% ACN, 0.1% formic acid), in a 90 min linear gradient. MS .raw data were acquired using the XCalibur software (Thermo Fisher Scientific). MS. raw files were processed using MaxQuant software^17^ (version 1.6.3.3) and searched against the human UniProt database (release 2016_07, 70,630 sequences) using the Andromeda search engine^18^ with the following settings: the parent mass and fragment ions were searched with an initial mass deviation of 4.5 p.p.m. and 20 p.p.m., respectively. Carbamidomethyl (C) was added as a fixed modification and Acetyl (N-term) and Oxidation (M) as variable modifications. The minimum peptide length was set to seven amino acids and a maximum of two missed cleavages and specificity for trypsin cleavage were required. The false discovery rates (FDR) at the protein and peptide level were set to 1%. The label-free quantification (LFQ) setting was enabled for protein quantification^19^. Razor and unique peptides were used for quantification. Perseus software^20^ (version 1.6.2.2) was used for statistical analysis. Data were filtered to remove potential contaminants, reverse peptides which match a decoy database and peptides only identified in their modified form. LFQ intensities were transformed by log_2_. A 2 sample t-test was used to determine significantly regulated proteins, with the permutation-based FDR ≤ 0.05 and S0 = 0.1 being considered significant.

### Atomic force microscopy

For imaging purposes, the ECMs prepared as above were fixed with 2%PFA and stored with PBS containing 1% Penicillin/Streptomycin at 4°C. A day prior to imaging, the dishes were washed with distilled water 5 times to wash off any salt and then air dried overnight. Samples were imaged by intermittent contact mode in air using a Bruker ScanAsyst. The probe was auto-tuned using Nanoscope software. Images were taken at 10×10 μm and 2×2 μm area at least two sites. Data were processed using Nanoscope analysis software 1.4 prior to image export. The Roughness (Rq) values were determined using the software. Roughness is the root mean square average of the image and is calculated based on the height difference per pixel along the sample length. Rq is used to study the surface topography of various nanostructures^21,22^. Rq provides a quantitative measure of fibril organisation in dermis and could possibly suggest the integrity of matrix^23^.

### Collagen degradation assay

To quantify the degradation of collagen in different fibroblast cell lines a collagen degradation assay based on^24^ was used. DQ Collagen, type I from bovine skin, Fluorescein conjugate (Invitrogen) was used to coat the wells of a 96-well plate for one hour at 37°C and quenched with 20mM glycine for 5 minutes. Next, fibroblasts were plated at a concentration of 2.5 x10^3^/well and incubated in normal culture conditions for 18 hours. Cells were then fixed with 4% paraformaldehyde for 20 minutes at room temperature, permeabilised with Triton 0.1% X-100 for 5 minutes at room temperature and stained with Alex Fluor 546 Phalloidin (Invitrogen) 1:2000 for one hour and DAPI (Invitrogen) 1:2000 for 15 minutes. Wells were imaged with the Opera Phenix High Content Screening System (PerkinElmer, Inc.), and collagen degradation was quantified by either measuring the area where DQ-collagen had been degraded and normalising to cell number (DAPI) using a custom ImageJ Macro or by scoring of images by 2 independent assessors (one of them blinded), scoring the intensity of collagen degradation as low (1: peri-cytoplasmic ring of collagen degradation less than 1/4 of the cytoplasmic diameter); medium (2: peri-cytoplasmic ring of collagen degradation approximately 1/3 of the cytoplasmic diameter) or high (3: peri-cytoplasmic ring of collagen degradation approximately 1/2 of the cytoplasmic diameter) (Extended Data Fig. 3a). To establish the ratio of degradation between shCtrl-HFF and shCtrl-UV-HFF, and *shMMP1*-HFF and *shMMP1*-UV-HFF cells, we first generated a score (H) of collagen degradation for each condition: Σ(number of images of collagen degrading fibroblasts of low intensity*1)+(number of images of collagen degrading fibroblasts of medium intensity*2)+(number of images of collagen degrading fibroblasts of high intensity*3)/total number of images. We scored, 150 images for shCtrl-HFF, 147 images for shCtrl-UV-HFF, 150 images for *shMMP1*-HFF, and 146 images for *shMMP1*-UV-HFF fibroblasts. The ratios were established as: Ratio shCtrl-UV-HFF/ shCtrl-HFF = H-score shCtrl-UV-HFF/ H-score shCtrl-HFF, and Ratio *shMMP1*-UV-HFF/*shMMP1*-HFF = H-score *shMMP1*-UV-HFF / H-score *shMMP1*-HFF.

### MMP1 ELISA

MMP1 in the secretome of cell lines was quantified with a MMP1 Human ELISA Kit (Thermo Fisher Scientific) according to manufacturer’s protocol. Secretomes collected from cells were diluted 1:10 and standards and samples were measured in triplicate. Samples were incubated on plate overnight at 4°C. Absorbance measured on a Spectra Max M5 plate reader (Molecular Devices).

### Quantitative PCR

RNA was collected in duplicate from 1×10^6^ cells lysed in trizol (company) after secretomes were collected. The aqueous phase of phenol-chloroform separation was collected and RNA extracted using RNeasy Blood and Tissue Kit (Qiagen). Concentration was determined with the Qubit RNA HS Assay (Invitrogen) and 500ng RNA was reverse transcribed to cDNA using TaqMan Reverse Transcription Reagents (Thermo Fisher) and diluted 1:20 in nuclease free water. Genes were quantified by qPCR using TaqMan Gene expression assays and Fast Mastermix on a QuantStudio 3 system. *GAPDH* (Hs02758991_g1) and *ACTB* (Hs01060665_g1) were used as housekeeping genes. *MMP1* (Hs00899658_m1), *COL1A1* (Hs00164004_m1), and *COL1A2* (Hs01028956_m1) were quantified and normalised to the geometric mean of both housekeeping genes and relative expression calculated using 2^−Δct^.

### Western Blots

Intracellular protein was extracted from cells using the NucBuster Protein Extraction Kit (Merck) and quantified using Pierce BCA Protein Assay Kit (Thermo Fisher Scientific). 40μg of cytoplasmic protein was diluted in Laemmli Buffer (Bio-Rad) with Beta-mercaptoethanol, denatured at 95°C for 5 min and loaded onto Mini-PROTEAN TGX Gels (Bio-Rad). Samples were transferred to nitrocellulose membranes using the TransBlot Tubro system (Bio-Rad) and protein visualised with Ponceau Stain (G-Biosciences). Membranes were blocked in 5% BSA TBS-T (5% BSA in 1x Tris Buffered Saline, 0.1% Tween 20, Sigma Aldrich) for 1 hour and incubated with primary antibodies overnight in 5% BSA TBS-T (MMP1 1:1000 ab137332, MMP2 1:1000 D4M2N Cell Signalling, B-actin 1:10,000 ab8226, Abcam). Membranes were washed with TBS-T and incubated in secondary antibodies (Dnk pAb to Rb IgG IRDye 680RD, ab216779, Goat pAb to Ms IgG IRDye 800CW, ab216772, Abcam) for 1 hour at room temperature. Membranes were visualised using the Odyssey CLx system (Licor).

**Methods Figure 1.**
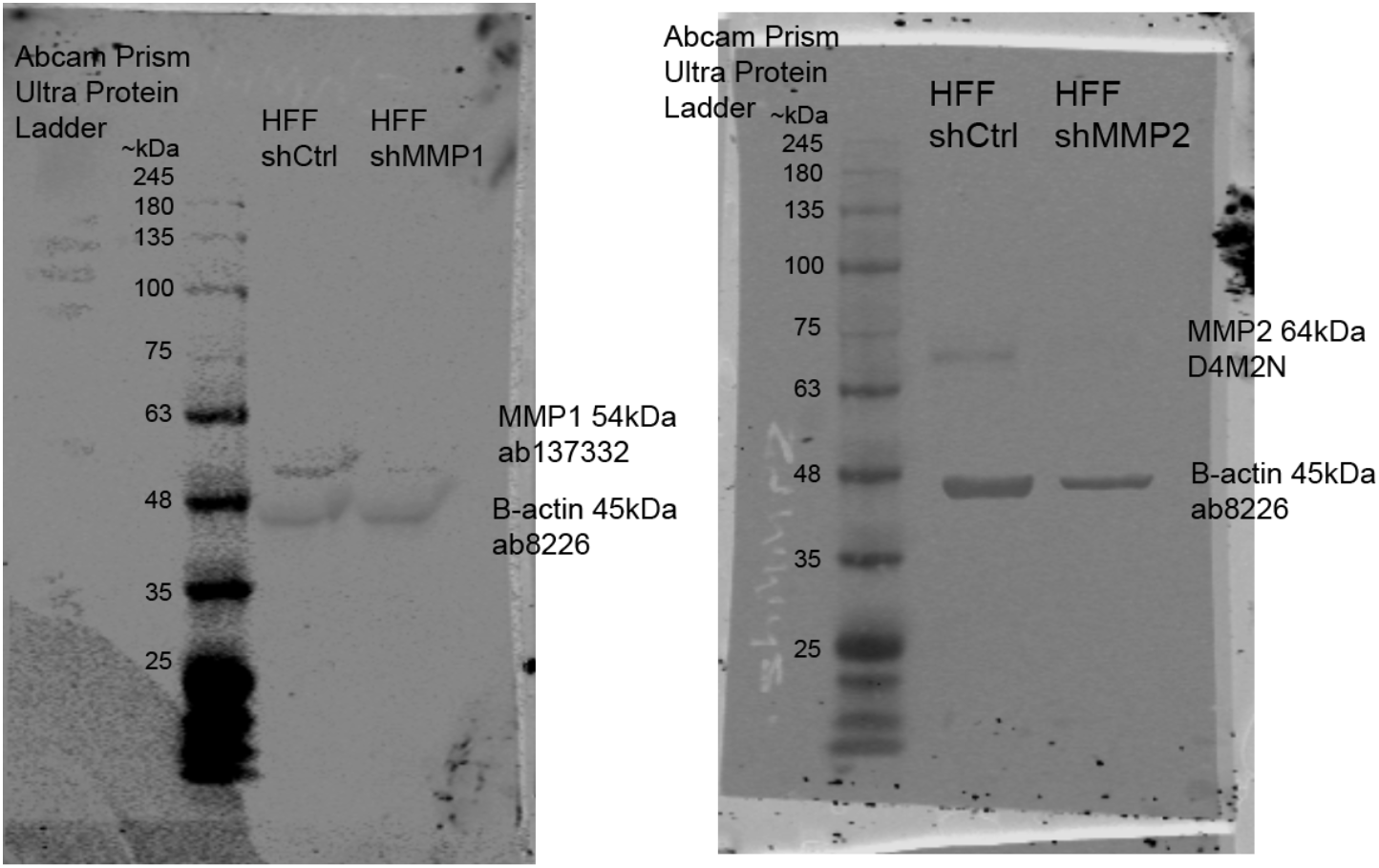
Uncropped whole images of western blots for MMP1, MMP2 and B-actin in shRNA knockdown HFF cell lines.

### Melanoma spheroid invasion assay

Melanoma cell lines were cultured in U-bottom 96-well plates (Brand) at 1×10^3^ cells per well, spun at 200xg, and spheroids allowed to form over 72 hours. Culture media was removed from the wells and 100ul collagen (PureCol, Advanced BioMatrix) added to the wells. Plates were briefly spun for 15sec at 200xg and incubated at 37°C for 1hr to set collagen. For concentration gradient collagen was diluted to 0.25, 0.5, 1, 1.5, 2 or 2.5mg/ml in phenol free DMEM without FCS. For degraded collagen invasion collagen was diluted to 1.5mg/ml in phenol free DMEM containing Collagenase I (Gibco) to final concentration in the collagen of 0.5, 1, 5, or 10μg/ml. For invasion using fibroblast secretomes collagen was diluted to 1.5mg/ml using secretome collected from various fibroblast cell lines instead of DMEM. Spheroids were allowed to invade over 72 hours and light microscopy photographs were taken. Images were analysed to quantify the invasive area around the spheroid by creating a layered mask for the spheroid core and invasive area and quantifying the invasive area as a percentage of the total size of the spheroid in ImageJ software. For single cell invasion analysis spheroids in the above conditions were imaged using an Opera Phenix (Perkin Elmer) with a 5x (need NA) lens. Spheroids were stained with Hoechst 33342 for 1 hour prior to imaging. Stacks of images were acquired through the full depth of the spheroids, and images were analysed using Columbus software (Perkin Elmer). A custom analysis pipeline was used to detect individually invading cells. In brief, a maximal intensity projection of the Hoechst stain was processed (Flatfield correction: Basic, Guassian blur: 2px) was used to determine the perimeter of the spheroid, and nuclei were detected in the remaining portion of the image. Experiments with varying levels of collagen matrices or collagenase were repeated and confirmed by an independent laboratory, blinded for matrix composition.

### Organotypic 3D invasion models

Melanoma invasion through fibroblast modified collagen was assayed using a protocol adapted from ^25^. Briefly, equal numbers of HFF, UV-HFF, *shMMP1*-HFF, and *shMMP1*-UV-HFFfibroblasts were mixed with Collagen I, rat tail (Corning) and cultured in 35mm culture dishes. Collagen disks were allowed to contract until they fit in a 24 well plate. Cell suspensions of Sk-mel-28 and A375 at 4×10^4^ cells/ml were plated on top of each collagen disc in duplicate for each fibroblast condition. Cells and collagen were cultured as normal for approximately 5 days. Collagen discs were then transferred to Falcon 3.0μm high density PET membrane (Corning) in Falcon 6-well Deep Well TC-treated Polystyrene Plates (Corning) to create an air/liquid interface to drive melanoma invasion into collagen. After 10 days constructs were fixed in 4% paraformaldehyde and embedded in paraffin and stained with H&E and stained for Fibronectin with FN1 antibody (F3648, Sigma Aldrich). For each construct the number of cells invading into the collagen was counted in at least five different fields of view under light microscopy, by 2 independent scorers.

### Second Harmonic Generation Imaging

Second harmonic generation (SHG) imaging of collagen in 3D organotypic models was performed using a Leica SP8 upright confocal microscope with 25x NA 0.95 water objectives controlled by Leica LAS X software. The images were acquired with 880nm illumination laser line from MaiTai Ti:Sapphire laser (Spectra Physics) and HyD-RLD detector installed 440/20nm filter cube (Leica), also used 483/32nm filter (Leica) collecting autofluorescence signals at the same time. Z-series optical sections were collected with a step-size of 1 micron driven by SuperZ galvo stage (Leica). Collagen was quantified in ImageJ by measuring the mean signal intensity of z-stack sum projections in three equal sized areas acorss three fields of views for each collagen disc.

### Clinical samples

Three international patient cohorts of primary cutaneous melanoma were used in this paper. The A cohort (n=31), the B cohort (n=222) and the C cohort (n=113). All clinical and pathological information assessed was done under appropriate institutional ethics approvals. Comprehensive clinical outcome was available for the B and C cohorts; and was collected prospectively at both institutions. The correlation between solar elastosis and invasion of melanoma cells at the invasive front (IF) was done in the A cohort in patients with invasive primary melanoma where a distinct vertical growth was determined in patients aged ≥ 55 at the time of diagnosis. The correlation and histological assessments of the B and C cohorts were done in primary cutaneous melanomas of patients aged ≥ 55 at the time of diagnosis with Breslow ≥ 1*mm*. The clinical characteristics of the cohorts are described in Extended Data Table S5.

### Histological and clinical sample analysis

The histological assessment of primary cutaneous melanomas of the A, B and C (20%) cohorts, from three international centers, was performed by at least two observers (Cohort A: observer 1, 2; Cohort B: 1, 3, 4; Cohort C: 1, 2). Discrepancy in Cohorts A and B were jointly reviewed and consensus agreed. There was high interobserver agreement in cohort C (>0.65), and all scores were done blinded for clinical outcome. The survival analyses were performed by members of the team who did not score histological variables. We included all samples with sufficient material to assess the tumour body, invasive front (IF) and tumour-adjacent skin. Recurrent tumours were excluded. Non-primary melanomas were excluded. Solar elastosis was scored as described by Landi et al., and cutoffs for low, moderate and high solar elastosis; or CSD noCSD established from the original Landi categories^1^. Landi et al established a scoring system for the degree of solar elastosis from absent to severe using an 11-point score, from 0 to 3+. To generate binary categories, cases are classified as bearing chronic sun damage (CSD), for scores between 0 to 2-, or non-CSD for scores 2 to 3+. Cutoffs for absent (range 0, 0+), low (range 1-to 1+), moderate (range 2-to 2+) and high (3-to 3+) were established from the same range, as previously used^26^. We assessed the inter- reliability of the binary CSD classification between 2 scorers using the kappa statistic, which showed 0.75 concordance for the B cohort (weighted kappa= 0.75, 95% CI= 0.69-0.79).

The proportion of melanoma cell invasion at the IF in the dermis was scored in categories. We assessed the front of the melanoma component in the dermis that is in direct contact with the dermal matrix, and scored 0/1: no invasion/minimal invasion: <5% of cells in contact with the dermis are actively invading the matrix, detaching from the IF of the melanoma; 2: low invasion: 5-25% of melanoma cells at the IF detaching from the tumour body; 3: moderate invasion: 25-50% of the IF is actively detaching from the main VGP and entering deeper structures; 4: high invasion: the majority of cells at the IF are independently interacting with the matrix, detached from the body of the tumour. Binary categories were then generated with low invasion (scores 0-2) and high invasion (3-4), a decision taken before performing survival analyses. We assessed the inter- reliability of the invasion score binary classification between 2 scorers using the kappa statistic, which showed 0.7 concordance for the C cohort (weighted kappa= 0.7, 95% CI= 0.64-0.77).

The amount of collagen at the IF of the tumour and in tumour-adjacent skin was scored from H&E slides (C cohort) according to abundance of distinctly formed collagen bundles. Two independent pathologists examined collagen on H&E routine stained sections of normal skin surrounding the melanomas and the collagen adjacent/enveloping the invasive front of the tumour in the dermis at 100-200x magnification. The following scoring system was used: collagen absent or low (1): when fully formed collagen bundles were rare, and the visible collagen was distributed in haphazard smaller fragments or unidentifiable in an amorphous deposit of elastotic material. Low collagen (2): when well-defined, undulating fibres of normal dermal length collagen are scarce, and a pattern of elastotic (fragmented or aggregate) material predominates. Medium collagen (3): well-defined, undulating and organised fibres coexist with aggregate elastotic material. High collagen (4): well-defined fibres in organised disposition predominating, with minimal or absent elastotic material interspersed between the tight bundles. We provide representative images below. We assessed the inter- reliability of the collagen score classification between 2 scorers using the kappa statistic, which showed 0.78 concordance for the C cohort (weighted kappa= 0.78, 95% CI= 0.7-0.81).

A subset of samples in Cohort A (n=16), for which tissue was available, were stained with trichrome of Masson to quantify collagen. We were unable to score the B cohort IF due to Covid-19 international restrictions.

**Methods Figure 2.**
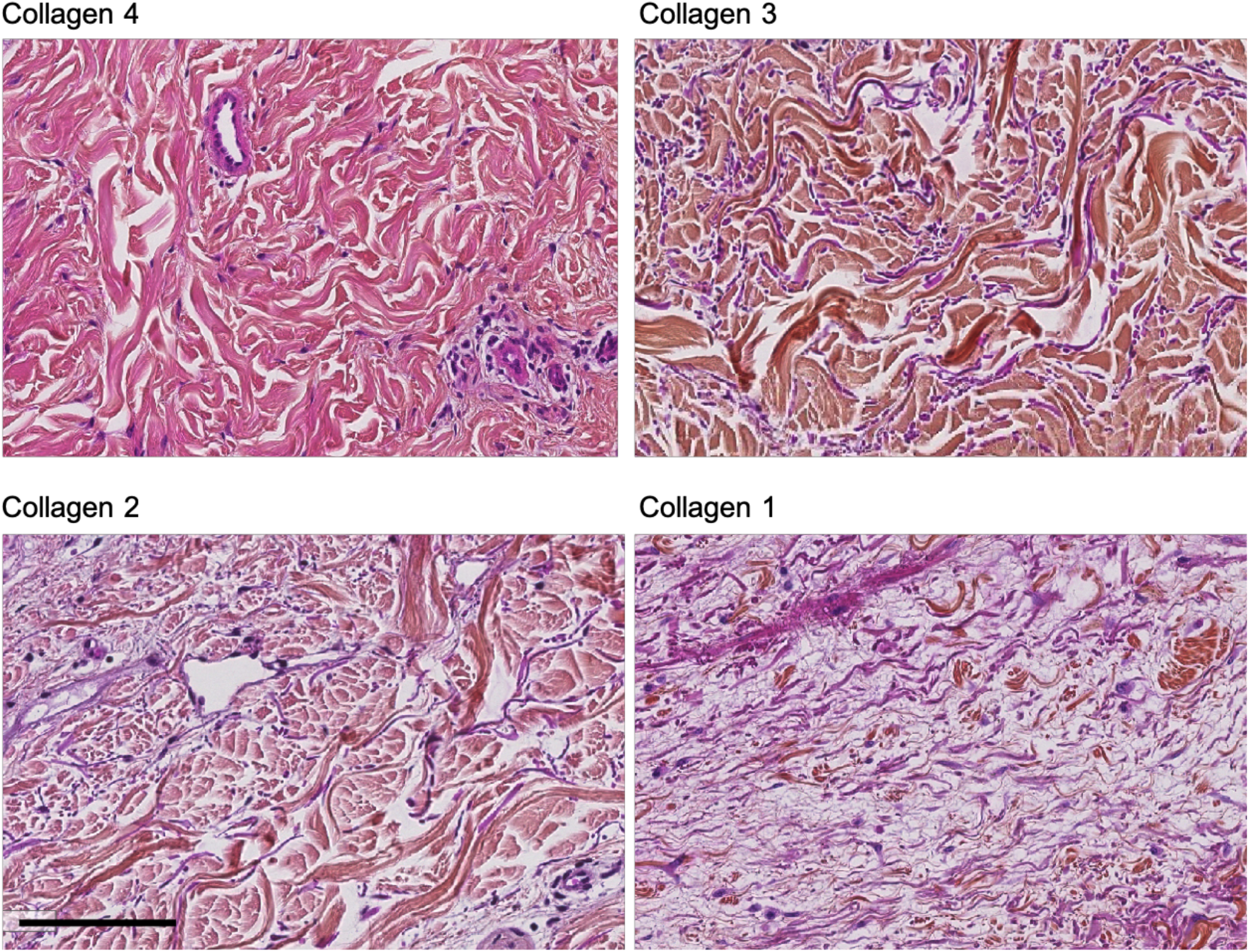
Representative photomicrographs of collagen scoring. 4: preservation of collagen fibres in the dermis. 3. Combination of preserved, normal collagen bundles (pink) intermixed with elastotic fibres of non-collagenous material (purple). 2. Combination of some collagen (pink) with purple elastotic fibres. Multiple fragmentation of collagen. 1. Scarce or absent collagen, substitution of the matrix by elastotic heterogeneous fibres (purple). Rare collagen fragments interspersed (pink).

### Statistical Analysis

For *in vitro* studies statistical analysis was performed in GraphPad Prism (version 7.01, GraphPad Software, Inc.). For comparisons between two groups Mann Whitney tests were used and for comparisons between multiple groups Kruskal-Wallis with Dunn’s multiple comparison tests. A p-value <0.05 was considered significant, after correcting for multiple testing where necessary. For human studies statistical analysis was performed in R (version 3.5.1, RStudio v1.2.5001, RStudio Inc). Association between categorical data was performed with Fisher exact tests. Survival analysis was performed using survival (version 3.1-12) and survminer (version 0.4.6) packages. For all clinical cohorts, melanoma specific survival (MSS), overall survival (OS) and progression free (PFS) were calculated from time of diagnosis. Univariate grouped survival analysis performed with Kaplan Meier and Log Rank tests, and multivariate analyses with cox regression models, with evaluation of the proportional hazard assumption. Gene expression (log_2_(x + 1) normalised RSEM) and clinical data from the TCGA SKCM and PANCAN data sets was accessed from the UCSC Xena data portal (https://xenabrowser.net/datapages/). Samples were grouped into COL1A1 high or low based on the expression relative to the median expression of all samples. The melanoma associated fibroblast (MAF) score for each sample was determined by calculating the geometric mean of all genes in a published melanoma associated fibroblast signature^27^. High and low MAF samples were classified based on the median signature score.

### Code Availability

No custom codes or algorithms were used in the analysis of this study and all analysis packages and software is cited in manuscript. R codes used are available from the corresponding author upon request.

## References

1. Gilchrest, B. A., Eller, M. S., Geller, A. C. & Yaar, M. The pathogenesis of melanoma induced by ultraviolet radiation. N Engl J Med 340, 1341–1348 (1999).

2. Whiteman, D. & Green, A. The pathogenesis of melanoma induced by ultraviolet radiation. N Engl J Med 341, 766–767 (1999).

3. Whiteman, D. C. et al. Melanocytic nevi, solar keratoses, and divergent pathways to cutaneous melanoma. J Natl Cancer Inst 95, 806–812 (2003).

4. Shain, A. H. & Bastian, B. C. From melanocytes to melanomas. Nat. Rev. Cancer 16, 345–358 (2016).

5. Saini, N. et al. The Impact of Environmental and Endogenous Damage on Somatic Mutation Load in Human Skin Fibroblasts. 1–25 (2016). doi:10.1371/journal.pgen.1006385

6. Tsai, S., Balch, C. & Lange, J. Epidemiology and treatment of melanoma in elderly patients. Nat. Rev. Clin. Oncol. 7, 148–152 (2010).

7. Balch, C. M. Decreased Survival Rates of Older-Aged Patients with Melanoma: Biological Differences or Undertreatment? Ann. Surg. Oncol. 22, 2101–2103 (2015).

8. Sahai, E. et al. A framework for advancing our understanding of cancer-associated fibroblasts. Nat. Rev. Cancer 20, 174–186 (2020).

9. Kalluri, R. The biology and function of fibroblasts in cancer. Nat. Rev. Cancer 16, 582–598 (2016).

10. Viros, A. et al. Ultraviolet radiation accelerates BRAF-driven melanomagenesis by targeting TP53. Nature 511, 478–482 (2014).

11. Berwick, M. et al. Sun Exposure and Mortality From Melanoma. J. Natl. Cancer Inst. 97, 195–199 (2005).

12. Mishra, K. et al. Histopathologic variables differentially affect melanoma survival by age at diagnosis. Pigment Cell Melanoma Res 32, 593‐600 (2019).

13. Howard, M. D. et al. Anatomic location of primary melanoma: Survival differences and sun exposure. J Am Acad Dermatol 81, 500–509 (2019).

14. Berwick, M. et al. Sun Exposure and Melanoma Survival: A GEM Study. Cancer Epidemiol Biomarkers Prev 23, 2145–2152 (2014).

15. Vollmer, R. T. Solar Elastosis in Cutaneous Melanoma. Am. J. Clin. Pathol. 128, 260–264 (2007).

16. Green, A., Baade, P., Coory, M., Aitken, J. & Smithers, M. Population-Based 20-Year Survival Among People Diagnosed With Thin Melanomas in Queensland, Australia. J. Clin. Oncol. 30, 1462‐1467 (2012).

17. National Cancer Institute. The Surveillance, Epidemiology, and End Results (SEER) Program. Available at: www.seer.cancer.gov. Accessed June 2019.

18. Cancer Research UK. https://www.cancerresearchuk.org/health-professional/cancer-statistics/statistics-by-cancer-type/melanoma-skin-cancer/mortality#heading-One

19. Hashim, D. et al. Cancer mortality in the oldest old: a global overview. Aging (Albany. NY). 12, 16744–16758 (2020).

20. Stanta, G., Campagner, L., Cavallieri, F. & Giarelli, L. Cancer of the oldest old. What we have learned from autopsy studies. Clin. Geriatr. Med. 13, 55–68 (1997).

21. Ahmadzadeh, H. et al. Modeling the two-way feedback between contractility and matrix realignment reveals a nonlinear mode of cancer cell invasion. Proc. Natl. Acad. Sci. U. S. A. 114, E1617–E1626 (2017).

22. Levental, K. R. et al. Matrix crosslinking forces tumor progression by enhancing integrin signaling. Cell 139, 891–906 (2009).

23. Arnold, S. A. et al. Lack of host SPARC enhances vascular function and tumor spread in an orthotopic murine model of pancreatic carcinoma. Dis. Model. Mech. 3, 57–72 (2010).

24. Driskell, R. R. et al. Distinct fibroblast lineages determine dermal architecture in skin development and repair. Nature 504, 277–281 (2013).

25. Kaisers, W. et al. Age, gender and UV-exposition related effects on gene expression in in vivo aged short term cultivated human dermal fibroblasts. PLoS One 12, 1–21 (2017).

26. Alexandrov, L. B. et al. Signatures of mutational processes in human cancer. Nature 500, 415–21 (2013).

27. Consortium, G. Genetic effects on gene expression across human tissues. (2017). doi:10.1038/nature24277

28. Fisher, G. et al. Molecular basis of sun-induced premature skin ageing and retinoid antagonism. Nature 335–9. (1996).

29. Fisher, G. J. et al. Pathophysiology of Premature Skin Aging Induced by Ultraviolet Light. N. Engl. J. Med. 337, 1419–1429 (1997).

30. Webb, H. K. et al. Roughness parameters for standard description of surface nanoarchitecture. Scanning 34, 257–263 (2012).

31. Girasole, M. et al. Roughness of the plasma membrane as an independent morphological parameter to study RBCs: a quantitative atomic force microscopy investigation. Biochim. Biophys. Acta 1768, 1268–1276 (2007).

32. Argyropoulos, A. J. et al. Alterations of Dermal Connective Tissue Collagen in Diabetes: Molecular Basis of Aged-Appearing Skin. PLoS One 11, e0153806 (2016).

33. Kligman AM. Early destructive effect of sunlight on human skin. JAMA 210, 2377–80 (1969).

34. Landi, M. T. et al. MC1R germline variants confer risk for BRAF-mutant melanoma. Science (80-.). 313, 521–522 (2006).

35. Vincent, K. & Postovit, L. Investigating the utility of human melanoma cell lines as tumour models. Oncotarget 8, 10498‐10509 (2017).

36. Nakyai, W., Saraphanchotiwitthaya, A., Viennet, C., Humbert, P. & Viyoch, J. An In Vitro Model for Fibroblast Photoaging Comparing Single and Repeated UVA Irradiations. Photochem. Photobiol. 93, 1462–1471 (2017).

37. Knott, A. et al. Decreased fibroblast contractile activity and reduced fibronectin expression are involved in skin photoaging. Journal of dermatological science 58, 75–77 (2010).

38. Schwartz, E., Feinberg, E., Lebwohl, M., Mariani, T. J. & Boyd, C. D. Ultraviolet radiation increases tropoelastin accumulation by a post-transcriptional mechanism in dermal fibroblasts. J. Invest. Dermatol. 105, 65–69 (1995).

39. Tirosh, I. et al. Dissecting the multicellular ecosystem of metastatic melanoma by single-cell RNA-seq. Science (80-.). 352, 189–196 (2016).

40. Kaur, A. et al. SFRP2 in the aged microenvironment drives melanoma metastasis and therapy resistance. Nature 532, 250–254 (2016).

41. Kaur, A. et al. Remodeling of the Collagen Matrix in Aging Skin Promotes Melanoma Metastasis and Affects Immune Cell Motility. Cancer Discov. 9, 64–81 (2019).

42. Kai, F., Drain, A. P. & Weaver, V. M. Review The Extracellular Matrix Modulates the Metastatic Journey. Dev. Cell 49, 332–346 (2019).

43. Lu, P., Weaver, V. M. & Werb, Z. The extracellular matrix: a dynamic niche in cancer progression. J. Cell Biol. 196, 395–406 (2012).

44. Trucco, L. D. et al. Ultraviolet radiation–induced DNA damage is prognostic for outcome in melanoma. Nat. Med. (2018). doi:10.1038/s41591-018-0265-6

45. Liu, D. et al. Integrative molecular and clinical modeling of clinical outcomes to PD1 blockade in patients with metastatic melanoma. Nat. Med. 25, 1916–1927 (2019).

46. Wolf, Y. et al. UVB-Induced Tumor Heterogeneity Diminishes Immune Response in Melanoma. Cell 179, 219–235 (2019).

47. Kuczek, D. E. et al. Collagen density regulates the activity of tumor-infiltrating T cells. J. Immunother. Cancer Vol. 7(1):68, (2019).

48. Pickup, M. W., Mouw, J. K. & Weaver, V. M. The extracellular matrix modulates the hallmarks of cancer. EMBO Rep. 15, 1243–1253 (2014).

49. Miskolczi, Z. et al. Collagen abundance controls melanoma phenotypes through lineage-specific microenvironment sensing. Oncogene 37, 3166–3182 (2018).

50. Condeelis, J. & Segall, J. E. Intravital imaging of cell movement in tumours. Nat. Rev. Cancer 3, 921–930 (2003).

51. Provenzano, P. P. et al. Collagen reorganization at the tumor-stromal interface facilitates local invasion. BMC Med. 4, 38 (2006).

52. Wang, W. et al. Single cell behavior in metastatic primary mammary tumors correlated with gene expression patterns revealed by molecular profiling. Cancer Res. 62, 6278–6288 (2002).

## References

1 Landi, M. T. et al. MC1R germline variants confer risk for BRAF-mutant melanoma. Science (New York, N.Y.) 313, 521–522, doi:10.1126/science.1127515 (2006).

2 Bolger, A. M., Lohse, M. & Usadel, B. Trimmomatic: a flexible trimmer for Illumina sequence data. Bioinformatics 30, 2114–2120, doi:10.1093/bioinformatics/btu170 (2014).

3 Li, H. & Durbin, R. Fast and accurate short read alignment with Burrows-Wheeler transform. Bioinformatics 25, 1754–1760, doi:10.1093/bioinformatics/btp324 (2009).

4 Koboldt, D. C. et al. VarScan 2: somatic mutation and copy number alteration discovery in cancer by exome sequencing. Genome Res 22, 568–576, doi:10.1101/gr.129684.111 (2012).

5 McLaren, W. et al. Deriving the consequences of genomic variants with the Ensembl API and SNP Effect Predictor. Bioinformatics 26, 2069–2070, doi:10.1093/bioinformatics/btq330 (2010).

6 Cornelius, L. A. et al. Selective upregulation of intercellular adhesion molecule (ICAM-1) by ultraviolet B in human dermal microvascular endothelial cells. J Invest Dermatol 103, 23–28, doi:10.1111/1523-1747.ep12388971 (1994).

7 Dobin, A. et al. STAR: ultrafast universal RNA-seq aligner. Bioinformatics 29, 15–21, doi:10.1093/bioinformatics/bts635 (2013).

8 Cibulskis, K. et al. Sensitive detection of somatic point mutations in impure and heterogeneous cancer samples. Nat Biotechnol 31, 213–219, doi:10.1038/nbt.2514 (2013).

9 Blokzijl, F., Janssen, R., van Boxtel, R. & Cuppen, E. MutationalPatterns: comprehensive genome-wide analysis of mutational processes. Genome Medicine 10, 33, doi:10.1186/s13073-018-0539-0 (2018).

10 Love, M. I., Huber, W. & Anders, S. Moderated estimation of fold change and dispersion for RNA-seq data with DESeq2. Genome Biology 15, 550, doi:10.1186/s13059-014-0550-8 (2014).

11 Jassal, B. et al. The reactome pathway knowledgebase. Nucleic Acids Res 48, D498–d503, doi:10.1093/nar/gkz1031 (2020).

12 Cukierman, E. Cell migration analyses within fibroblast-derived 3-D matrices. Methods Mol Biol 294, 79–93, doi:10.1385/1-59259-860-9:079 (2005).

13 Franco-Barraza, J., Beacham, D. A., Amatangelo, M. D. & Cukierman, E. Preparation of Extracellular Matrices Produced by Cultured and Primary Fibroblasts. Current protocols in cell biology 71, 10.19.11-10.19.34, doi:10.1002/cpcb.2 (2016).

14 Püspöki, Z., Storath, M., Sage, D. & Unser, M. Transforms and Operators for Directional Bioimage Analysis: A Survey. Advances in anatomy, embryology, and cell biology 219, 69–93, doi:10.1007/978-3-319-28549-8_3 (2016).

15 Hernandez-Fernaud, J. R. et al. Secreted CLIC3 drives cancer progression through its glutathione-dependent oxidoreductase activity. Nat Commun 8, 14206, doi:10.1038/ncomms14206 (2017).

16 Rappsilber, J., Ishihama, Y. & Mann, M. Stop and go extraction tips for matrix-assisted laser desorption/ionization, nanoelectrospray, and LC/MS sample pretreatment in proteomics. Anal Chem 75, 663–670, doi:10.1021/ac026117i (2003).

17 Cox, J. & Mann, M. MaxQuant enables high peptide identification rates, individualized p.p.b.-range mass accuracies and proteome-wide protein quantification. Nat Biotechnol 26, 1367–1372, doi:10.1038/nbt.1511 (2008).

18 Cox, J. et al. Andromeda: a peptide search engine integrated into the MaxQuant environment. J Proteome Res 10, 1794–1805, doi:10.1021/pr101065j (2011).

19 Cox, J. et al. Accurate Proteome-wide Label-free Quantification by Delayed Normalization and Maximal Peptide Ratio Extraction, Termed MaxLFQ. Mol Cell Proteomics 13, 2513–2526, doi:10.1074/mcp.M113.031591 (2014).

20 Tyanova, S. et al. The Perseus computational platform for comprehensive analysis of (prote)omics data. Nat Methods 13, 731–740, doi:10.1038/nmeth.3901 (2016).

21 Webb, H. K. et al. Roughness parameters for standard description of surface nanoarchitecture. Scanning 34, 257–263, doi:10.1002/sca.21002 (2012).

22 Girasole, M. et al. Roughness of the plasma membrane as an independent morphological parameter to study RBCs: a quantitative atomic force microscopy investigation. Biochimica et biophysica acta 1768, 1268–1276, doi:10.1016/j.bbamem.2007.01.014 (2007).

23 Argyropoulos, A. J. et al. Alterations of Dermal Connective Tissue Collagen in Diabetes: Molecular Basis of Aged-Appearing Skin. PloS one 11, e0153806, doi:10.1371/journal.pone.0153806 (2016).

24 Sabeh, F. et al. Tumor cell traffic through the extracellular matrix is controlled by the membrane-anchored collagenase MT1-MMP. J Cell Biol 167, 769–781, doi:10.1083/jcb.200408028 (2004).

25 Timpson, P. et al. Organotypic collagen I assay: a malleable platform to assess cell behaviour in a 3-dimensional context. J Vis Exp, e3089, doi:10.3791/3089 (2011).

26 Moreno, A. et al. Histologic Features Associated With an Invasive Component in Lentigo Maligna Lesions. JAMA dermatology 155, 782–788, doi:10.1001/jamadermatol.2019.0467 (2019).

27 Tirosh, I. et al. Dissecting the multicellular ecosystem of metastatic melanoma by single-cell RNA-seq. Science (New York, N.Y.) 352, 189–196, doi:10.1126/science.aad0501 (2016).

